# RABEP1 amplifies front signaling in neutrophil migration

**DOI:** 10.1101/2025.05.08.652481

**Authors:** Daniel H. Kim, Ramizah Syahirah, Conwy Zheng, Chang Ding, Alan Y. Hsu, Sheng Liu, Jun Wan, Qing Deng

## Abstract

Neutrophils are the first responders of our innate immune system, crucial for defense against various infections. The intricate regulation of neutrophil migration is essential for neutrophil function. However, a complete mechanistic understanding is missing. We performed a miRNA overexpression screen and identified miR-190 as a potent suppressor of neutrophil migration in zebrafish. Through a second round of small-scale screening using neutrophil-specific knockouts of putative miR-190 targets, we identified that *rabep1* (Rabaptin-5, RAB GTPase binding effector protein 1) is essential for neutrophil motility and chemotaxis in zebrafish. Re-expressing full-length *RABEP1* in the knockout, but not a truncated form lacking the Rab4/Rab5 binding domain, rescued cell motility. Knocking down *RABEP1* in human dHL-60 cells consistently reduced cell motility. *RABEP1*-deficient dHL-60 cells accumulate excessive members inside the cells. The Rab5 GTP level is unaffected, but the *RABEP1* knockdown cells displayed reduced PAK phosphorylation and overall reduced actin polymerization, but still appropriately polarized upon chemokine stimulation. Overexpression of dominant-negative Rab4 or Rab5 similarly inhibits neutrophil migration. Our data suggests that *RABEP1* drives endosomal recycling, Rac activation, and leading-edge actin polymerization, providing significant insights into the role of the endocytic pathway in neutrophil motility.

## INTRODUCTION

Neutrophils are essential innate immune cells that respond to infections and inflammation.^1–2^ Neutrophils migrate to the inflammation sites, expressing and releasing cytokines to recruit other cells of the immune system while also directly attacking foreign microorganisms through degranulation, phagocytosis, production of reactive oxygen species (ROS), and generation of neutrophil extracellular traps.^3–5^ However, deregulated neutrophil activity can lead to various diseases, such as inflammatory diseases, autoimmune diseases, or cancer.^6–8^ Therefore, the regulation of neutrophil migration is essential to maintain a balanced and efficient immune system.^9–11^ The specific molecules involved in the modulation of neutrophil activity or the exact mechanisms behind such regulations are not yet well understood. MiRs are short, conserved, noncoding RNAs that function as transcriptome regulators.^12^ They bind to their target genes’ 3’ untranslated regions and recruit the RNA-induced silencing complex, downregulating their expression.^13–14^ Until now, most miR studies have been conducted using transgenic mouse models with only a handful focused on neutrophil activity, which entails limitations such as difficulty visualizing the effects of miR on neutrophil motility *in vivo* or non-conclusive results due to non-specific inflammation induced by invasive methods for the imaging of neutrophil activity.^15–16^ To overcome such limitations, we have utilized the zebrafish model, which shares a highly similar immune system with mammals. It also supports noninvasive live imaging since zebrafish are transparent through their embryonic and larval stages. ^17–20^

Our previous study screening a wide array of miRs identified nine hits significantly decreasing neutrophil chemotaxis when overexpressed in zebrafish neutrophils.^21^ We verified this initial observation using two different *miR-190* OE founder fishlines. Subsequently, we performed transient knockout of putative *miR-190* target genes, and the Rab GTPase Binding Effector Protein 1 (*rabep1*) gene showed the highest significance in neutrophil motility change compared to the control. ^22^

*Rabep1* is the protein-coding gene for Rabaptin-5, an effector protein for Rab4 and Rab5.^23^ Members of the Rab family, including Rab4 and Rab5, have been identified as key regulators of vesicular trafficking, which is crucial for regulating cell migration through activating signaling cascades or by internalizing and recycling receptors.^24–26^ However, whether endosomal recycling regulates neutrophil migration has not been determined. To this end, we investigated the phenotype of *RABEP1* knockdown in differentiated human leukemia (dHL-60) cells. The *RABEP1*-deficient cells were defective in motility, with excessive internalized endosomes and reduced Rac activation, indicating that *RABEP1* is essential for endosome recycling and precise regulation of cell migration in neutrophils.

## MATERIALS AND METHODS

### Reagents

ATCC Universal Mycoplasma Detection Kit (ATCC, 50-238-3275), HBSS (Gibco, 14025-092), Puromycin (Gibco, A1113803), Penicillin-Streptomycin (Gibco, 15140122), GlutaMax (Gibco, 35050-0611), OPTI-MEM (Gibco, 51985-034), pHrodo™ Green *E. coli* BioParticles™ Conjugate (Invitrogen, P35366), Amplex Red Hydrogen Peroxide Peroxidase Assay Kit (Invitrogen, A22188), eBioscience™ Flow Cytometry Staining Buffer (Invitrogen, 00-4222-26), Novex™ Tris-Glycine SDS Running Buffer (Invitrogen, LC26754), Pierce™ 10X Western Blot Transfer Buffer (35040), PageRuler™ Prestained Protein Ladder (26616), RIPA Lysis and Extraction Buffer (89900), Pierce™ Protease Inhibitor Tablets (A32965), Lipofectamin 3000 Transfection Kit (L3000015), PVDF Transfer Membranes (88518), 10 x PBS (BP3994), Sudan Black (BP109-10), DMSO (327182500), TEMED (BP150-20), Ammonium Persulfate (17874), Agarose (BP160-500), 20 x TBS Tween20 (28360), Tween20 (AAJ20605AP), Triton X-100 (AAA16046AP), Bovine Serum Albumin (BP1600-100), KOH (437135000), NaCl (BP358-1), Sodium Phosphate (BP332-500), KCl (BP366-500), CaCl_2_ (C79-500), MgSO_4_ (BP213-1) and H_2_O_2_ (H325-500) were from ThermoFisher Scientific (Waltham, MA, USA). dPBS (21-031-CV), RPMI-1640 (10-041-CV), DMEM (10-013-CV), HEPES (25-060-CI), Falcon® 12-well Clear Flat Bottom TC-treated Multiwell Cell Culture Plate (353043), Corning® 100 mm Not TC-treated Culture Dish (430591), Falcon® 100 mm TC-treated Cell Culture Dish (353003), Falcon® 96-well Cell Clear Flat Bottom TC-treated Culture Microplate (353075), Sodium Bicarbonate (25-035CI) and Sodium Pyruvate (25-000-CI) were from Corning (Corning, NY, USA). MISSION® TRC2 pLKO.5-puro Non-Mammalian shRNA Control Plasmid DNA (SHC202), MISSION^®^ pLKO.1-puro-CMV-TurboGFP^™^ Positive Control Plasmid DNA (SHC003), MISSION® TRC2 pLKO.5-puro *RABEP1* shRNA 1 Plasmid DNA (TRCN0000327713), MISSION® TRC2 pLKO.5-puro *RABEP1* shRNA 2 Plasmid DNA (TRCN0000363576), MISSION® TRC2 pLKO.5-puro *RABEP1* shRNA 3 Plasmid DNA (TRCN0000369958), glucose (G8270-100G), Fibrinogen (F3879), polybrene (TR-1003-G), PMA (P1585), TCA (T6399), NaF (G9422), β-glycerophosphate (S6776) and fMLP (F3506) were from Millipore Sigma (Burlington, MA, USA). 30% Acrylamide/Bis Solution (1610156), 10% SDS Solution (1610416), 2× Laemmli sample buffer (1610737), Resolving Gel Buffer (1610798) and Stacking Gel Buffer (1610799) were from Bio-Rad Laboratories (Hercules, CA, USA). PrimeSTAR MAX DNA Polymerase (R045), In-Fusion® Snap Assembly Master Mix (ST2320), SOC media (ST0215), Lenti-X concentrator (631232) and Stellar™ Competent Cells (636763) were from Takara Bio (San Jose, CA, USA). Plasmids pCMV-dR8.2 dvpr (8455), pCMV-VSV-G (8454), Rab5DN (35139), Rab5 CA (35140), TagRFP-T-EEA1 (42635) were from Addgene (Watertown, MA, USA). Geneblocks for Rab4 DN and Rab11 DN were from Twist Biosciences (San Francisco, CA, USA). Versaladder DNA ladder (D012-500) and Low Melt Agarose (A-204-25) were from GoldBio (St. Louis, MO, USA). Ibidi μ-Slide 8 Well High ibiTreat (80806) and µ-Slide 8 Well High Glass Bottom (80807) were from IBIDI (Fitchburg, WI, USA). Leukotriene B_4_ (20110) was from Cayman Chemicals (Ann Arbor, MI, USA). HyClone™ Characterized Fetal Bovine Serum (SH30396.03) was from Cytiva (Marlborough, MA, USA). VECTASHIELD HardSet Antifade Mounting Medium (H-1400-10) was from VectorLabs (Newark, CA, USA). Paraformaldehyde Solution (15710) was from Electron Microscopy Sciences (Hatfield, PA, USA). Human *RABEP1* cDNA Clone (HG19536-ACGLN) was from Sino Biological (Wayne, PA, USA). Rab5 Pull-Down Activation Assay Kit (83701) was from NewEast Biosciences (King of Prussia, PA, USA). Black Wall 96-well Plate (655096) was from Greiner Bio-One (Monroe, NC, USA). Ethanol 200 Proof (2701) was from Decon Labs (Bryn Mawr, PA, USA). Precision Red Advanced Protein Assay (ADV02) was from Cytoskeleton, Inc. (Denver, CO, USA). Antibodies for immunoblotting were anti-*RABEP1* (HPA019669-25ul), anti-Vinculin (V9264), and anti-α-Tubulin (T5168) from Millipore Sigma (Burlington, MA, USA). Phospho-PAK1 (Ser199/204)/PAK2 (Ser192/197) Antibody (2605S) and PAK2 (3B5) Mouse mAb (4825S) from Cell Signaling (Danvers, MA, USA); secondary antibodies used were: Goat anti-Mouse IgG (H+L) Secondary Antibody (DyLight™ 680) (Invitrogen, 35518), Goat anti-Rabbit IgG (H+L) Secondary Antibody (DyLight™ 800 4X PEG) (Invitrogen, SA535571) from ThermoFisher Scientific (Waltham, MA, USA). Antibodies for immunofluorescence were Wheat Germ Agglutinin (Invitrogen, W32465), Alexa Fluor™ 488 Phalloidin (Invitrogen, A12379), and DAPI (Invitrogen, D3571) from ThermoFisher Scientific (Waltham, MA, USA). Phospho-Myosin Light Chain 2 (Ser19) Mouse mAb (3675) from Cell Signaling (Danvers, MA, USA); secondary antibodies used were: Goat anti-Mouse IgG (H+L) Secondary Antibody, DyLight™ 680 (Invitrogen, 35518). Antibodies for flow cytometry were BD Horizon™ BV421 Annexin V (BDB563973) and anti-Mouse IgM, κ Isotype Control (BDB562704) from BD Biosciences (San Jose, CA, USA). Alexa Fluor® 647 anti-human CD11b Antibody (301319) and Alexa Fluor® 647 Mouse IgG1, κ Isotype Ctrl (FC) Antibody (400130) from Biolegend (San Diego, CA, USA). Propidium Iodide (P3566) from ThermoFisher Scientific (Waltham, MA, USA). Primers were from Integrated DNA Technologies (Coralville, IA, USA).

### Animals

The zebrafish (*Danio rerio*) experiments were conducted in accordance with internationally accepted standards. The Animal Care and Use Protocols were approved by the Purdue Animal Care and Use Committee (PACUC), adhering to the Guidelines for Use of Zebrafish in the NIH Intramural Research Program (Protocol number: 1401001018).

Transient neutrophil-specific knockout zebrafish were generated by injecting Tol2 backbone plasmids into LyzC-Cas9-expressing embryos at their one-cell stage.

Plasmids for transient neutrophil-specific knockout of different genes were constructed using the following primers:

lats1 guide1 F: GGAGGGCCGAGAAACCCGAGTTTAAGAGCTATGCTGGAAACAGCATAGC

lats1 guide1 R: CCTGGAAGTTTATCCTGTTCGAACTAGGAGCCTGGAGAACTGC

lats1 guide2 F: GGATAAACTTCCAGGGTTTAAGAGCTATGCTGGAAACAGCATAGC

lats1 guide2 R: GTTTCTCGGCCCTCCCGAACCAAGAGCTGGAGGGAGAGGCTGTAT

gcat guide1 F: GGTGTTTTACAGCCCAGCAGTTTAAGAGCTATGCTGGAAACAGCATAGC

gcat guide1 R: CGGCGCGGATAGAGTCCAGCGAACTAGGAGCCTGGAGAACTGC

gcat guide2 F: ACTCTATCCGCGCCGGTTTAAGAGCTATGCTGGAAACAGCATAGC

gcat guide2 R: GGGCTGTAAAACACCCGAACCAAGAGCTGGAGGGAGAGGCTGTAT

rabep1-1 guide1 F: GAGCTGTCCGGGCGGCCAGGTTTAAGAGCTATGCTGGAAACAGCATAGC

rabep1-1 guide1 R: ATACAGCTTACCTGCCCCCCGAACTAGGAGCCTGGAGAACTGC

rabep1-1 guide2 F: GCAGGTAAGCTGTATGTTTAAGAGCTATGCTGGAAACAGCATAGC

rabep1-1 guide2 R: CCGCCCGGACAGCTCCGAACCAAGAGCTGGAGGGAGAGGCTGTAT

rabep1-2 guide1 F: CCAACCGGTATCCAGATGGGTTTAAGAGCTATGCTGGAAACAGCATAGC

rabep1-2 guide1 R: TCCGAAGACCATCCTTATACGAACTAGGAGCCTGGAGAACTGC

rabep1-2 guide2 F: AGGATGGTCTTCGGAGTTTAAGAGCTATGCTGGAAACAGCATAGC

rabep1-2 guide2 R: CTGGATACCGGTTGGCGAACCAAGAGCTGGAGGGAGA

arhgap20 guide1 F: GAAGTTACATTGGAGGTCGGTTTAAGAGCTATGCTGGAAACAGCATAGC

arhgap20 guide1 R: TGCGTCAGCCTCGATCCCTCGAACTAGGAGCCTGGAGAACTGC

arhgap20 guide2 F: ATCGAGGCTGACGCAGTTTAAGAGCTATGCTGGAAACAGCATAGC

arhgap20 guide2 R: CTCCAATGTAACTTCCGAACCAAGAGCTGGAGGGAGAGGCTGTAT

vamp8 guide1 F: GCACGAGCGACCTTCTGGGGTTTAAGAGCTATGCTGGAAACAGCATAGC

vamp8 guide1 R: CGGGCCAGAATTCGATCAACGAACTAGGAGCCTGGAGAACTGC

vamp8 guide2 F: TCGAATTCTGGCCCGGTTTAAGAGCTATGCTGGAAACAGCATAGC

vamp8 guide2 R: GAAGGTCGCTCGTGCCGAACCAAGAGCTGGAGGGAGA

praf2 guide1 F: AGTAATCCGACGATTCCGGGTTTAAGAGCTATGCTGGAAACAGCATAGC

praf2 guide1 R: ATGGGCCTTGTACTGGAAGCGAACTAGGAGCCTGGAGAACTGC

praf2 guide2 F: CAGTACAAGGCCCATGTTTAAGAGCTATGCTGGAAACAGCATAGC

praf2 guide2 R: AATCGTCGGATTACTCGAACTAGGAGCCTGGAGAACTGC

Transgenic zebrafish lines were generated by co-injecting Tol2 backbone plasmids with Tol2 transposase mRNA into wild-type AB embryos at their one-cell stage.

Plasmids for *miR-190* overexpression were constructed using the following primers:

miR-190a F: CTGGCAGTACGGGCTGCCTCTCCTCCTCTACCCAT

miR-190a R: TAACAGCAGTTGGCTCAAGCACAGTGGGCCTAAGA

Plasmids for neutrophil-specific knockout of *rabep1* were constructed using the following primers:

rabep1-1 guide1 F: GAGCTGTCCGGGCGGCCAGGTTTAAGAGCTATGCTGGAAACAGCATAGC

rabep1-1 guide1 R: ATACAGCTTACCTGCCCCCCGAACTAGGAGCCTGGAGAACTGC rabep1-1 guide2 F: GCAGGTAAGCTGTATGTTTAAGAGCTATGCTGGAAACAGCATAGC

rabep1-1 guide2 R: CCGCCCGGACAGCTCCGAACCAAGAGCTGGAGGGAGAGGCTGTAT

Plasmids for *RABEP1* rescue were constructed using the following primers:

RFP-CAAX linearization F: CTAACATGCGGTGACGTGGAGGAGAATCCCGGCCCTATGGTGTCTAAGGGCGAAGAG CTG

RFP-CAAX linearization R: GGTGGCGAGGTACCTGTATCACTG

RABEP1 linearization F: AGGTACCTCGCCACCATGGCGCAGCCGGGC

RABEP1 linearization R: GTCACCGCATGTTAGAAGACTTCCTCTGCCCTCTGTCTCAGGAAGCTGGTTAATGTCT GT

RABEP1 Δ5-2 F: AGGCCTCAAAAGATCAGGAGGATGATGAACAAC

RABEP1 Δ5-2 R: GATCTTTTGAGGCCTCCAGCTCTTTAATTTTG

### Cell Culture

HEK293T (CRL-11268) and HL-60 (CCL-240) were from the American Type Culture Collection (ATCC, Manassas, VA, USA). All cells were maintained at 37°C with 5% CO_2_ in a Forma™ Steri-Cycle™ i160 CO2 Incubator (NC1207547, ThermoFisher Scientific). HL-60 cells were cultured in RPMI-1640 supplemented with 10% FBS, 25 mM HEPES, 1% penicillin-streptomycin, 1% sodium bicarbonate, 1% sodium pyruvate, and 1% GlutaMax. HEK293T cells were cultured in DMEM supplemented with 10% FBS and 1% sodium bicarbonate. HL-60 cells were differentiated with 1.3% DMSO for 6 days. Cells were checked monthly for Mycoplasma using the ATCC Universal Mycoplasma Detection Kit.

To generate *RABEP1* knockdown HL-60 cell lines pLKO.5 lentiviral constructs with shRNA (*RABEP1* shRNA 1: TRCN0000327713, *RABEP1* shRNA 2: TRCN0000363576, *RABEP1* shRNA 3: TRCN0000369958) were used, and MISSION® TRC2 pLKO.5-puro Non-Mammalian shRNA Control Plasmid DNA was used as a non-targeting control.

The *RABEP1* rescue lentiviral constructs were generated by replacing TurboGFP of the *RABEP1* shRNA 3 inserted into MISSION^®^ pLKO.1-puro-CMV-TurboGFP^™^ Positive Control Plasmid DNA backbone with shRNA resistant full-length *RABEP1* or Δ5-2 truncated *RABEP1* using the following primers:

shRNA – turboGFP linearization F: AATTCTCGACCTCGAGACAAATGGC

shRNA – turboGFP linearization R: GGTGGCGACCGGGAGCGC

Rabep1 insert linearization F: CTCCCGGTCGCCACCATGGCGCAGCCGGGCCCG

Rabep1 insert linearization R: TCGAGGTCGAGAATTTCAGGAGAGCACACACTTGC

shRNA 369958 loop swap F: CATTACTCGAGTAATGATTCAGTTCTTTAACCTTTTTGAATTCAGTTATTAATAGT

shRNA 369958 loop swap R: ATTACTCGAGTAATGATTCAGTTCTTTAACCCCGGTGTTTCGTCCTTTCCA

shRNA 369958 resistant F: TAAAGGAGCTTAACCACTATCTGGAAGCTGAGAAATCTTGT

shRNA 369958 resistant R: GGTTAAGCTCCTTTACCTTTGAGGCCTCCAGCTCT

Lentiviral constructs with pCMV-dR8.2 dvpr and pCMV-VSV-G were co-transfected into HEK293T cells using the Lipofectamin 3000 transfection kit to produce lentivirus. The virus supernatant was collected at 48 and 72 hpt and then concentrated with a Lenti-X concentrator. HL-60 cells were spin-infected (2500g, 1.5 hrs) with 3 mL of complete RPMI-1640 medium containing concentrated lentivirus supplemented by 4 µg/ml polybrene. Infected HL-60 cells were then selected with 2 µg/ml puromycin to generate stable lines.

### RNA Sequencing

RNA sequencing was performed as described previously. ^21^ Briefly, whole kidney marrows of *Tg(LyzC-miR-190-dendra2)^pu38^* and *Tg(LyzC-vector-dendra2)^pu7^* were collected and processed into single-cell suspensions. Neutrophils were harvested and sorted by fluorescence-activated cell sorting (FACS). Total RNA was extracted and sent for RNA sequencing, which was performed at the Center for Medical Genomics at Indiana University School of Medicine. Samples were polyA enriched and sequenced with Illumina HiSeq 4000 ultra-low with reads ranging from 37M to 44M. The RNA-seq aligner from the STAR (v2.5) was employed to map RNA-seq reads to the reference genome, zebrafish (GRCz11), with previously the following parameter: “–outSAMmapqUnique 60”.^27^ Uniquely mapped sequencing reads were assigned to genes using featureCounts (from subread v1.5.1) with the following parameters: “-p -Q 10”.^28^ The functional analysis was performed on DEGs with a cutoff of FDR < 0.05 to identify significantly over-represented Gene Ontology (GO) and KEGG pathways by using the DAVID.^29^ The genes were filtered by counts per million, fold-change, *miR-190* target score, and p-values for further analysis. Finally, filtered zebrafish genes that did not have orthologues to humans were removed. RNA sequencing data have been uploaded online and are available at https://www.ncbi.nlm.nih.gov/geo/query/acc.cgi?acc=GSE167554, with the GEO accession number GSE167554.

### Microinjection

Microinjections of fish embryos were performed by injecting 1 nl of a mixture containing 25 ng/µl plasmid with or without 35 ng/µl Tol2 transposase mRNA into the cytoplasm of embryos at the one-cell stage, depending on the transient or transgenic purpose.

### Tailfin Wounding

Tailfin wounding was carried out on 3dpf zebrafish larvae, where the tailfin was cut as close to the CHT as possible and incubated at 28 °C for 1 hr. After incubation, the larvae were fixed with 4% paraformaldehyde for 2 - 24 hrs in preparation for Sudan Black staining.

### LTB4 Bathing

LTB4 chemoattractant bathing was carried out on 3 dpf zebrafish larvae, where the larvae were subjected to 30 nM LTB4 at 28 °C for 15 min. After incubation, the larvae were fixed with 4% paraformaldehyde for 2 - 24 hrs in preparation for Sudan Black staining.

### Whole Body Neutrophil Counting

Whole-body neutrophil counting was carried out on 3 dpf zebrafish larvae, which were either imaged under an AXIO Zoom V16 microscope (Zeiss, Thornwood, NY, USA) for GFP signal or fixed with 4% paraformaldehyde for 2 - 24 hrs in preparation for Sudan Black staining.

### Sudan Black Staining

Sudan Black staining was used to stain neutrophils after larval fixation. Staining was done for 30 min and then washed with 70% EtOH. The stained larvae were rehydrated with PBST (0.1% Tween-20) and subjected to pigment clearing by bathing in 1% KOH and 1% H_2_O_2_ at RT for 10 min. After depigmentation, the sample was washed three times with PBS and stored at 4°C for future analysis.

### Neutrophil Motility Assay

Neutrophil motility assay was carried out on 3-dpf zebrafish larvae, where the larvae were anesthetized with tricaine methanesulfonate and loaded into 1.5% low-melt agarose. The larvae were placed into Ibidi μ-Slide 8 Well High ibiTreat and immediately settled to the bottom of the well. Neutrophil migration in the head of the larvae was recorded using a Lionheart FX Automated Microscope (Biotek, Winooski, VT, USA) with a 10× phase objective at 28 °C for 30 min with a 1 min interval. Representative time-lapse fluorescence images for neutrophil motility were acquired using an LSM 710 laser scanning confocal microscope (Zeiss, Thornwood, NY, USA) with a 20× objective lens at 28 °C for 30 min with a 1 min interval. The neutrophil movement was tracked using the ImageJ plugin MTrackJ.

### Western Blotting

dHL-60 cells were collected and pelleted by centrifugation at 250 x g and 4°C for 5 min and washed with dPBS once. Cells were resuspended in 2x protease inhibitor in RIPA buffer for lysing and incubated on ice for 20 min. Lysed cells were centrifuged at max speed, and the supernatant was transferred to a new tube. 10μl of protein was added into 1 ml of Precision Red Advanced Protein Assay solution, mixed by inverting 10 times and incubated at RT for 1 min. The mixture was transferred to a cuvette, and OD was measured using a BioPhotometer (Eppendorf, Hamburg, Germany), where 10 x OD read was calculated as the amount of protein (μg/µl). 20 μg of protein was mixed with RIPA buffer for a total volume of 20 µl for each sample and was boiled at 95°C for 15 min after adding 4 µl of 6 x SDS dye. Samples were then cooled on ice and centrifuged at max speed for 1 min before loading onto polyacrylamide gel (10%). All 24 µl of the sample was loaded into each well, and PageRuler™ Prestained Protein Ladder was used to indicate protein size. The gel was first run at 90V for 30 min, then at 140V for 1 hr in Novex™ Tris-Glycine SDS Running Buffer. After running, the gel was trimmed to remove the stacking gel portion and placed in a gel holder cassette as a sandwich of sponge - paper - gel - PVDF Transfer Membrane - paper - sponge. The transfer was run at 60V for 1.5 hrs in Pierce™ 10X Western Blot Transfer Buffer. After the transfer was completed, PVDF Transfer Membrane was cut to the size of the gel and blocked with 2.5% milk in PBST at RT for 1 hr. The blocked membrane was probed with primary antibodies diluted to 1:1000 in 1% BSA in PBST at 4°C overnight on a shaker. The primary antibody probed membrane was washed 3 times with PBST at 5 min intervals and probed with secondary antibodies diluted to 1:5000 in 1% BSA in PBST at RT for 30 min in the dark. The membrane was washed 3 times with PBST at 5 min intervals and subjected to imaging under an LI-COR Odyssey Gel Imaging Scanner (BioAgilytix, Durham, NC, USA). Protein preparation for Phospho-PAK western blot was done by first counting dHL-60 cells and calculating 5 × 10^6 cells/ml, which were then starved in serum-free RPMI for 1 hr. Cells were then aliquoted into 6 time-points of 0 sec, 30 sec, 1 min, 2 min, 5 min, and 10 min, which were stimulated with 100nM final concentration of fMLP for each time-point. Stimulation was stopped by adding ice-cold stop solution (20% TCA, 40 mM NaF, and 20 mM β-glycerophosphate) at a 1:1 ratio, then put on ice for 1 hr for lysing. Lysates were pelleted at 14,000 × g for 10 min, washed once with 1 ml of ice-cold 0.5% TCA, and resuspended in 2× Laemmli sample buffer. After protein preparation, a phospho-PAK western blot was done using the method described above.

### Rab5-GTP Pull-Down

A Rab5 Pull-Down Activation Assay Kit (NewEast Biosciences) was used to isolate active Rab5 from whole-cell lysate by affinity precipitation. dHL-60 cells were counted and resuspended in ice-cold lysis buffer at a concentration of 10^7^ cells/ml. Cells were then lysed by pipetting and centrifuged at 12,000 x g and 4°C for 10 minutes to clear. The supernatant was collected for immediate use. The collected cell lysate was then divided into two tubes, where one was used for obtaining the lysate Rab5 protein amount, and the other was used for affinity precipitation of active Rab5. Affinity precipitation was done by adding 1μl of anti-Rab5-GTP antibody and 20μl of resuspended bead slurry. This was left to incubate at 4°C for 1 hr. The beads were then centrifuged at 5,000 x g for 1 min. Pelleted beads were washed 3 times using lysis buffer, resuspended in 2× Laemmli sample buffer, and then boiled at 95°C for 5 min. The sample was prepared for immunoblotting by centrifuging at 5,000 x g for 10 sec. 15μl of pull-down supernatant and 15μl of cell lysate were used for western blotting. Anti-Rab5 Rabbit Polyclonal Antibody diluted to 1:500 in 3% BSA in TBST was used as primary antibody, and Goat anti-Rabbit IgG (H+L) Secondary Antibody (DyLight™ 800 4X PEG) diluted to 1:1000 in 3% BSA in TBST was used as secondary antibody.

### Under-Agarose Random Migration Assay

Ibidi μ-Slide 8 Well High ibiTreat or µ-Slide 8 Well high Glass Bottom slides were first coated by adding 500μl of fibrinogen at a concentration of 100nM and incubated for 1 hr at 37°C. Residual fibrinogen solution was aspirated, and the slides were washed with dPBS once, then subjected to blocking with 2% BSA in dPBS at 37°C for 30 min. Slides were rewashed with dPBS and left to air-dry overnight in the cell culture hood without UV. dHL-60 cells were collected and resuspended in mHBSS at 5 × 10^5 cells/ml. 5μl of these cells were added to the center of each well of the fibrinogen-coated slides and were topped with 500μl of 1.5% Low Melt Agarose in mHBSS. The slides were centrifuged at 2000 x g and 25°C for 5 min to allow cells to settle under the agarose gel. Imaging was done using a Lionheart FX Automated Microscope (Biotek, Winooski, VT, USA) with a 10× phase objective at 37 °C for 2 hrs with a 1 min interval. dHL-60 cell movement was tracked using the ImageJ plugin MTrackJ.

### Flow Cytometry Analysis

dHL-60 cells were collected and resuspended in 100µl mHBSS at a concentration of 1 × 10^^7^ cells/ml. Cells were stained with BD Horizon™ BV421 Annexin V, Alexa Fluor® 647 anti-human CD11b Antibody, and Propidium Iodide. Another aliquot of cells at the same concentration was stained with anti-Mouse IgM, κ Isotype Control, Alexa Fluor® 647 Mouse IgG1, κ Isotype Ctrl (FC) Antibody, and Propidium Iodide as the control group. All samples were incubated on ice for 30 min, centrifuged at 300 x g at 4°C for 5 min, then washed with eBioscience™ Flow Cytometry (FACS) Staining Buffer. Cells were centrifuged at the same parameters, then resuspended in 350 μl of FACS Staining Buffer and placed back on ice for flow cytometry analysis. Flow cytometry was performed using LSR Fortessa™ X-20 Cell Analyzer (BD Biosciences, Franklin Lakes, NJ, USA).

### Phagocytosis Assay

pHrodo™ Green E. coli BioParticles™ Conjugates (Invitrogen) were reconstituted in mHBSS and sonicated at 10% amplitude for 5 min with 10-sec intervals. 5 × 10^5 dHL-60 cells were resuspended in 100μl of BioParticle suspension and incubated at 37°C for 1 hr. Phagocytosis activity was stopped by placing the cells on ice, and 250 μL mHBSS was added to increase the total volume for flow cytometry analysis. A total number of 5 × 105 dHL-60 cells in 350 μl mHBSS was used as a control. Flow cytometry was performed using LSR Fortessa™ X-20 Cell Analyzer (BD Biosciences, Franklin Lakes, NJ, USA) to quantify geometric mean FITC intensity in the dHL-60 cell population.

### Reactive Oxygen Species Assay

Amplex Red Hydrogen Peroxide Peroxidase Assay Kit (Invitrogen) was used to detect ROS production after fMLP stimulation. dHL-60 cells were collected and resuspended in a 1 mL Krebs-Ringer phosphate (KRPG) solution at a concentration of 7.5 × 10^5 cells/mL. The KRPG solution consists of 145 mM NaCl, 5.7 mM sodium phosphate, 4.86 mM KCl, 0.54 mM CaCl_2_, 1.22 mM MgSO_4,_ and 5.5 mM glucose with a pH of 7.35. 20μl of cell suspension was added into each well of a Black Wall 96-well Plate along with 20μl of prepared H_2_O_2_ standards at concentrations of 0 - 90μM. The reaction mix was prepared by adding 60μl of 50μM Amplex Red and 120 μl of 10 U/ml Horseradish Peroxidase (HRP) in 12 mL KRPG solution. This reaction mix was further modified to have 100 nM fMLP or the same volume of DMSO added as a control. 100 μl of modified reaction mix was added to cells for three technical repeats, where 100 μl of non-modified reaction mix was added to H_2_O_2_ standards for a final concentration of 0 – 15μM. ROS readings were taken every 15 min for 45 min immediately after stimulation using a Synergy Neo2 (BioTek) at an excitation/emission wavelength of 490/550 nm.

### Immunofluorescence Staining

dHL-60 cells were resuspended in mHBSS and attached to fibrinogen-coated slides for 30 min. Cells were stimulated with 100 nM fMLP for 30 min and fixed with 3% paraformaldehyde in dPBS for 15 min at 37°C. Cells were then permeabilized in PBS with 0.1% Triton X-100 and 3% BSA at RT for 1 hr. Permeabilized cells were incubated with Wheat Germ Agglutinin, Alexa Fluor™ 488 Phalloidin, or Phospho-Myosin Light Chain 2 diluted to 1:100 in 3% BSA at 4°C overnight, depending on the staining purpose. The cells were stained with secondary antibodies diluted to 1:500 in 3% BSA and DAPI at RT for 1 hr. Stained cells were imaged for fluorescence using a Zeiss LSM 880 Confocal Laser Scanning Microscope (Zeiss, Thornwood, NY, USA) with a 63× oil immersion objective lens.

### Statistical Analysis

Statistical analysis was performed with Prism 6 (GraphPad), where the Mann-Whitney test was used to compare the two groups. Dunnett’s multiple comparisons test was used to compare more than three groups to determine statistical significance. Individual p-values are indicated in the figures. Each statistical analysis was done on all biological repeats, with different repeats indicated with varying shades of color in the resulting graphs.

## RESULTS

### Overexpression of *miR-190* in zebrafish neutrophils suppresses neutrophil random migration and chemotaxis

Our previous study discovered that the overexpression (OE) of miR-190s in zebrafish neutrophils decreases neutrophil recruitment to an ear infection or tail wound site.^21^ To validate the results, we comprehensively characterized the transgenic lines *Tg(LyzC-miR-190-dendra2)^pu38^*and *Tg(LyzC-vector-dendra2)^pu7^* that express miR-190 from the spliced intron or the Dendra2 Vector control-(Fig. 1A). Tailfin transection was performed to examine the recruitment of neutrophils to a wound site (Fig. 1B). The average number of neutrophils migrated the wound site within the first hour is significantly lower in the *miR-190* OE zebrafish compared to vector control zebrafish (Fig. 1C-D). We also used LTB4 as a chemoattractant to recruit neutrophils to the caudal fin from the caudal hematopoietic tissue (CHT) (Fig. 1E). The percentage of neutrophils that migrated out was significantly reduced in *miR-190* OE zebrafish compared to the vector control (Fig. 1F, G). Live imaging was performed at the zebrafish head mesenchyme to examine neutrophil motility, where neutrophils spontaneously move (Fig.1H; Movie 1). The *miR-190* OE neutrophils exhibit a significantly decreased speed compared to vector control (Fig.1I). Finally, to rule out the possibility of *miR-190* OE influencing neutrophil development or survival, total neutrophil numbers were quantified, which did not differ between vector control and *miR-190* OE fish (Fig.1J, K). All assays were done on offspring from two different *miR-190* OE founders to rule out artifacts associated with the transgene insertion sites (Fig.S1; Movie 2). Taken together, *miR-190* suppresses neutrophil motility and chemotaxis in all conditions examined.

**Figure 1.**
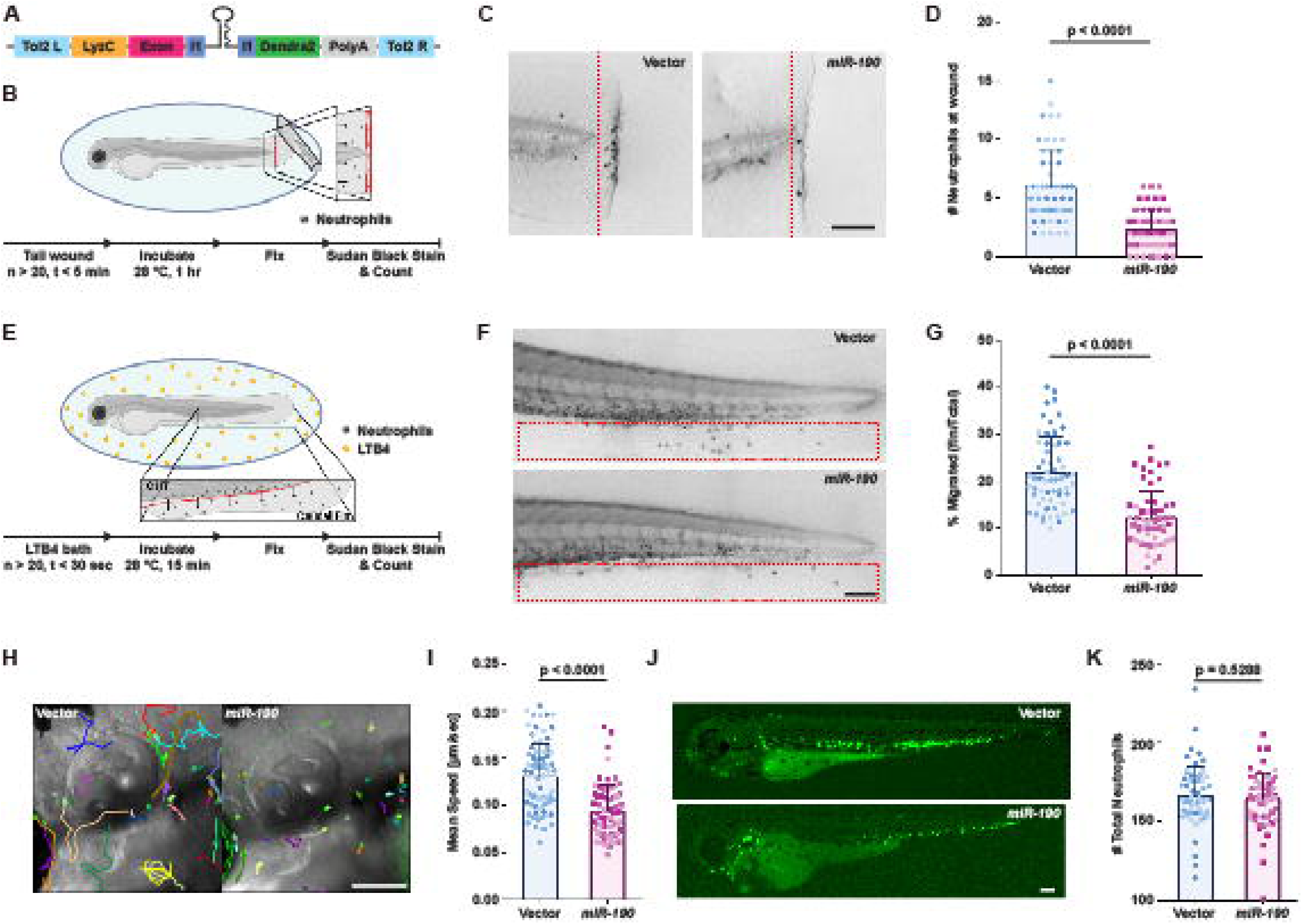
Overexpression of *miR-190* in zebrafish neutrophils leads to decreased cell motility and chemotaxis. (A) Schematic of Tol2-LyzC-miR-Dendra2 plasmid, which was injected into wild-type AB zebrafish embryos to generate *Tg(lyzC-miR-190-dendra2)^pu38^*and *Tg(lyzC-vector-dendra2)^pu7^* using the Tol2 transposon system. (B) Schematic of zebrafish tail wounding assay depicting Sudan black staining of wounded zebrafish to quantify the number of neutrophils that migrate to the wound within a specified time post-wounding. (C, D) Representative images and quantification of neutrophil recruitment. (Scale bar, 100μm) The assay had 3 biological repeats, each containing 17-20 fish per group. Quantification is presented as mean ± SD, using the Mann-Whitney test. (E) Schematic of LTB4 chemotaxis assay depicting Sudan black staining of zebrafish to quantify the percentage of neutrophils that migrate out of the CHT to the caudal fin within a specified time of LTB4 exposure. (F, G) Representative images and quantification of neutrophil chemotaxis percentage from CHT to caudal fin in vector and *miR-190* OE larvae with 15min LTB4 exposure. (Scale bar, 100μm) The assay had 3 biological repeats, each containing 20 fish per group. Quantification is presented as mean ± SD, using the Mann-Whitney test. (H, I) Representative images and quantification of live neutrophil motility imaging in vector and *miR-190* OE larvae for 30 min. (Scale bar, 100μm) The assay was done with 3 biological repeats, each containing 22-39 neutrophils tracked from 1-4 fish per group. Quantification is presented as mean ± SD, using the Mann-Whitney test. (J, K) Representative GFP images and Sudan black staining quantification of whole-body neutrophil count in vector and *miR-190* OE larvae. (Scale bar, 100μm) The assay was done with 3 biological repeats, each containing 20 fish per group. Quantification is presented as mean ± SD, using the Mann-Whitney test.

### A neutrophil-specific knockout screen identifies *rabep1* as a potential regulator for neutrophil migration in zebrafish

To understand how miR-190 regulates neutrophil migration, we sorted neutrophils from the whole kidney marrow, performed RNA sequencing, and identified differentially expressed genes (DEGs) upon miR-190 overexpression. GO enrichment analysis shows that the down-regulated DEGs represent biological processes involved with estrogen-dependent gene expression, chromatin organization, Rho GTPase effectors and signaling by Rho GTPases (Fig. 2B). To get to the direct target of miR-190, we further filtered the DEGs with *miR-190* target score (<-0.25), p-Value (<0.009) and fold-change (log_2_FC<-1.8), resulted in a list of 6 possible direct targets of *miR-190* (Fig. 2A; Table 1). We moved forward to transiently knock out each specifically in neutrophils using the CRISPR-Cas9 system to determine if any play a role in regulating neutrophil migration. Injecting a plasmid to express sgRNAs ubiquitously and GFP only in neutrophils into the neutrophil-specific Cas9 expressing line with an eye marker (*Tg(LyzC: Cas9, Cry: GFP/RFP)^pu34/pu52^*) allows tracking neutrophil motility without generating stable lines (Fig. 2C). Neutrophils with praf2 or *rabep1* gene knockout have a significantly decreased speed (Fig. 2D). To rule out possible *rabep1-1* sgRNA off-target effects, we used different sgRNAs (*rabep1-2*) to knockout *rabep1*. We observed the same phenotype, indicating that *rabep1* is required for neutrophil motility.

**Figure 2.**
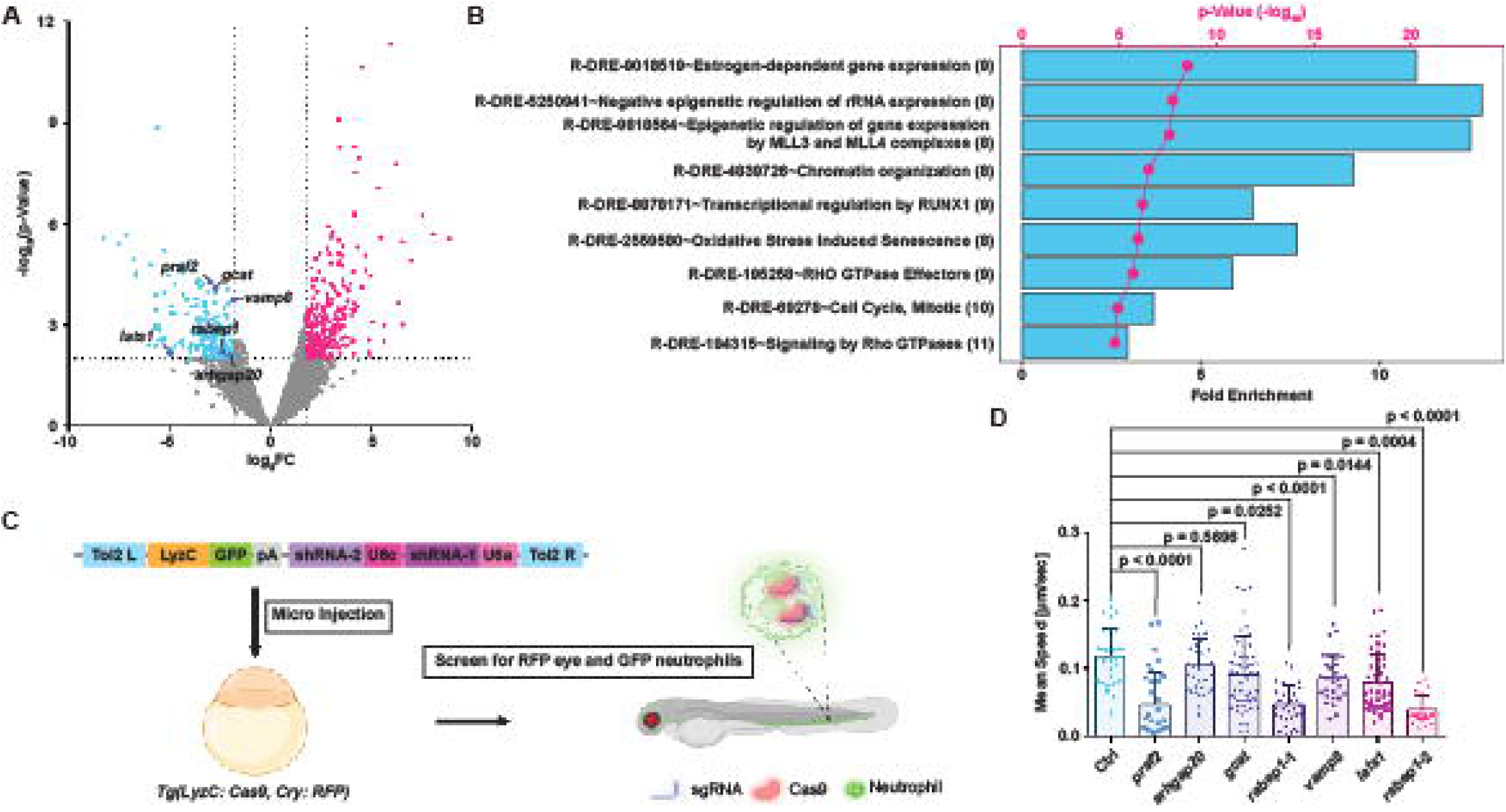
*miR-190* overexpression significantly downregulates *rabep1* expression in neutrophils. (A) Volcano plot of DEGs with significant expression changes in *miR-190* OE neutrophils compared with vector control. Down-regulated DEGs are shown in cyan, and up-regulated DEGs are shown in pink. Down-regulated DEGs selected through screening criteria are depicted in the plot. (B) Schematic of transient injection of guide-expressing plasmid into Cas9-expressing embryos to generate transient TSKO. sgRNA-expressing plasmids were designed to express two separate sgRNAs targeting one specific gene at two locations. (C) Quantification of live neutrophil motility imaging in transient TSKO compared to control TSKO. The assay was done with 3 biological repeats, each containing 20-58 neutrophils tracked from 5-8 fish per group. Quantification is presented as mean ± SD, using Dunnett’s multiple comparisons test. (D) Schematic of developing transgenic TSKO fishlines through crossing transgenic neutrophils specific Cas9 expressing fish with transgenic neutrophils specific guide RNA expressing fish.

**Table 1.** The short list of putative *miR-190* target genes. Gene name, accession number, *miR-190* target score, fold-change, and p-Value of the possible miR-190 direct targets in zebrafish neutrophils.

| Name | Gene Name | Accession Number | MiR 190 Target Score | log<sub>2</sub>FC | -log<sub>10</sub>(p-Value) |
| --- | --- | --- | --- | --- | --- |
| *praf2* | PRA1 domain family, member 2 | NM_007213 | -0.34 | -2.987 | 4.264 |
| *arhgap20* | Rho GTPase activating protein 20 | NM_020809 | -0.33 | -1.968 | 2.092 |
| *gcat* | Glycine C-acetyltransferase | NM_001172554.1 | -0.35 | -2.690 | 4.080 |
| *rabep1* | Rabaptin, RAB GTPase binding effector protein 1 | NM_004703 | -0.32 | -2.334 | 2.199 |
| *vamp8* | Vesicle-associated membrane protein 8 | NM_003761 | -0.30 | -1.818 | 3.758 |
| *lats1* | Large tumor suppressor kinase 1 | NM_004690 | -0.25 | -4.976 | 2.169 |

### *Rabep1* is essential for neutrophil random migration and chemotaxis in zebrafish

To further confirm the results of our transient screen, we generated lines with a neutrophil-specific knockout of *rabep1*(Fig.3A). Similar to *miR-190* OE, *rabep1* knockout resulted in reduced neutrophil recruitment to a wound site (Fig. 3B, C), impaired LTB4-induced mobilization (Fig. 3D, E), reduced motility in the head mesenchyme (Fig. 3F, G; Movie 3), without affecting whole-body neutrophil counts (Fig. 3H, I). The *RABEP1* gene codes for the Rabaptin-5 protein, an effector of small GTPase Rab4 and Rab5, facilitating membrane recycling or endosomal maturation of early endosomes.^23–26^ The human *RABEP1* gene is highly conserved with the zebrafish *rabep1* gene, sharing 70.94% homology. Rabaptin-5 has multiple Rab4 or Rab5 binding domains, which bind active Rab4 or Rab5, and a Rabex5 (Rab5 guanine nucleotide exchange factor) binding site, which facilitates activation of Rab5.^30–31^ A previous study characterized the functional domains of RABEP1 by rescuing its knockdown using various truncations. A truncation that deletes a Rab5 binding domain and part of a Rab4 binding domain (Δ5-2 *RABEP1*) exhibited the most robust phenotype with large and malformed endosome structures.^24^ Following the same strategy, we generated *Tg(LyzC:FL RABEP1-RFP-Caax)^pu53^* and *Tg(LyzC:* Δ*5-2 RABEP1-RFP-Caax)^pu54^* lines and crossed them with *rabep1* TSKO fish. A cross between *rabep1* TSKO and *Tg(LyzC: RFP-Caax)^pu54^* was used as a control. Full length *RABEP1,* but not Δ5-2 *RABEP1* expression, rescued the *rabep1* TSKO phenotype (Fig3.J-L; Movie 4).

**Figure 3.**
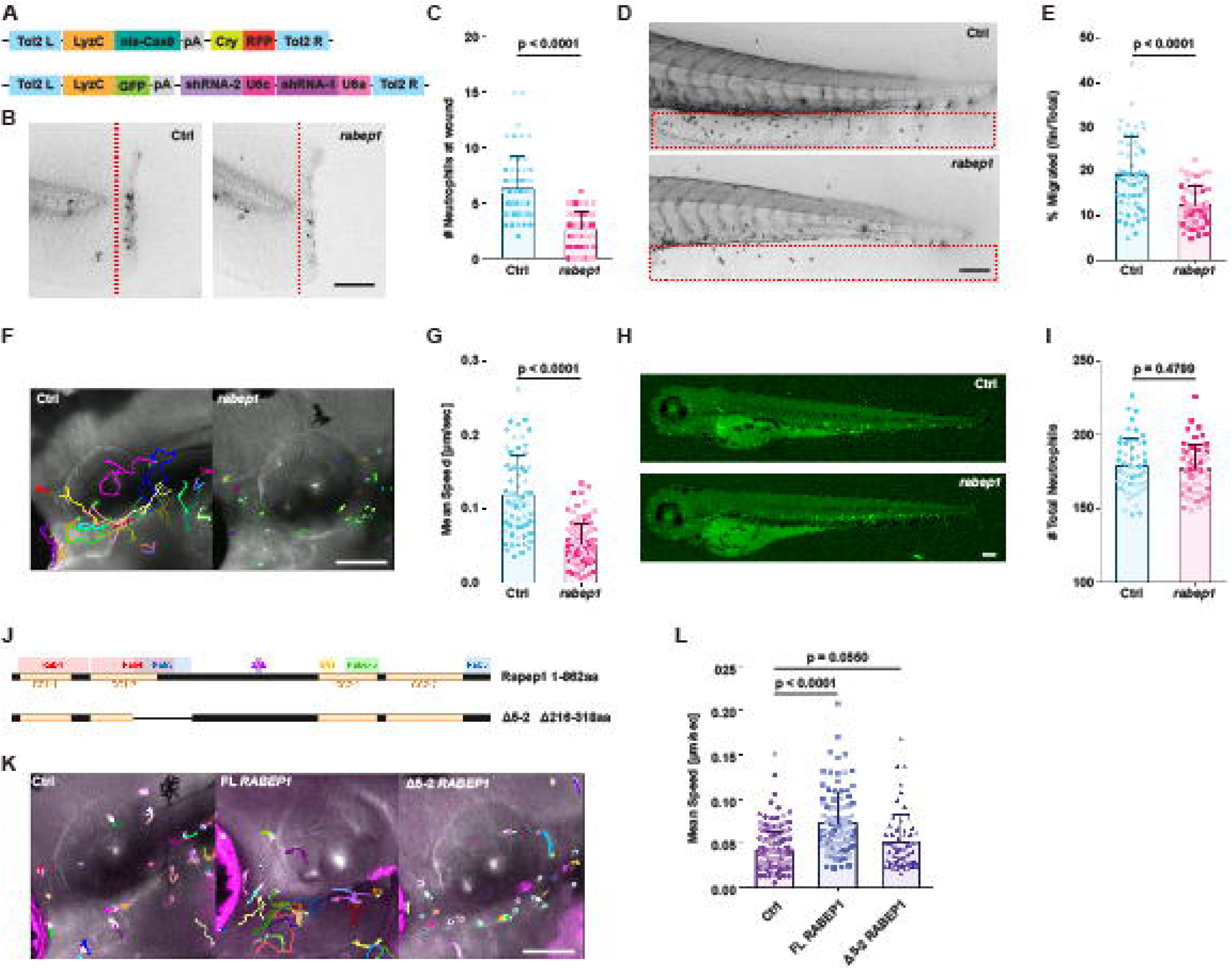
Knockout of *rabep1* in zebrafish neutrophils leads to decreased cell motility and chemotaxis. (A, B) Representative images and quantification of neutrophil recruitment to the tail wound site in *Tg(LyzC: ctrl sgRNAs, LyzC: Cas9, Cry: RFP)* and *Tg(LyzC: rabep1 sgRNAs, LyzC: Cas9, Cry: RFP)* larvae 1hr post wounding. (Scale bar, 100μm) The assay was done with 3 biological repeats, each containing 11-22 fish per group. Quantification is presented as mean + SD, using the Mann-Whitney test. (C, D) Representative images and quantification of neutrophil chemotaxis percentage from CHT to caudal fin in control TSKO and *rabep1* TSKO larvae with 15min LTB4 exposure. (Scale bar, 100μm) The assay was done with 3 biological repeats, each containing 16-20 fish per group. Quantification is presented as mean ± SD, using the Mann-Whitney test. (E, F) Representative images and quantification of live neutrophil motility imaging in control TSKO and *rabep1* TSKO larvae for 30 min. (Scale bar, 100μm) The assay was done with 3 biological repeats, each containing a total number of 19-38 neutrophils tracked from 3-4 fish per group. Quantification is presented as mean + SD, using the Mann-Whitney test. (G, H) Representative GFP images and Sudan black staining quantification of whole-body neutrophil count in control TSKO and *rabep1* TSKO larvae. (Scale bar, 100μm) The assay was done with 3 biological repeats, each containing 16-19 fish per group. Quantification is presented as mean ± SD, using the Mann-Whitney test. (I) Schematic of the full-length *RABEP1* protein, Rabaptin-5, and Δ5-2 *RABEP1* protein with domains annotated. (J, K) Representative images and quantification of live neutrophil motility imaging in *rabep1* TSKO x Tol2-LyzC-RFP control, *rabep1* TSKO x FL *RABEP1* OE, and *rabep1* TSKO x Δ5-2 *RABEP1* OE larvae for 30 min. (Scale bar, 100μm) The assay was done with 3 biological repeats, each containing 17-37 neutrophils tracked from 2-4 fish per group. Quantification is presented as mean + SD, using Dunnett’s multiple comparisons test.

### *RABEP1* is essential for dHL-60 cell migration

HL-60 cells become neutrophil-like when differentiated and show characteristics such as migration, chemotaxis, ROS production, and phagocytosis.^32^ We generated stable *RABEP1* knockdown cell lines through lentiviral transduction of three different shRNAs and confirmed knockdown (Fig. 4A, B). A cell line expressing the non-targeting control shRNA was used as a control. In the following experiments, we moved forward with the two cell lines with higher knockdown (shRNA2 and shRNA3). To determine if *RABEP1* affects cell differentiation, we used flow cytometry analysis to measure the surface expression of CD11b, a maturation marker. *RABEP1* shRNA2 and shRNA3 lines displayed over 50% or 90% CD11b-positive populations. The cell population is healthy with low apoptosis (Fig. 4C). Subsequently, we conducted under-agarose random migration assays on fibrinogen-coated slides and tracked the movements of dHL-60 cells. Both *RABEP1* knockdown cell lines had a significantly decreased speed compared to the control (Fig.4D, E; Movie 5). We measured phagocytosis and ROS production to investigate whether *RABEP1* is involved in other neutrophil functions. The phagocytic capabilities or ROS production rate in *RABEP1* knockdown cell lines are comparable to the control (Fig. 4F, G), indicating that *RABEP1* is particularly required for cell motility. We then performed a rescue similar to the zebrafish model, stably expressing TurboGFP, FL *RABEP1*, or Δ5-2 *RABEP1* in the shRNA3 line. Ctrl shRNA cell lines expressing TurboGFP were generated as a control. Western blotting of these cell lines confirmed a robust knockdown of *RABEP1* and the OE of both FL *RABEP1* and Δ5-2 *RABEP1*(Fig.5 A-B). The lines are all similarly differentiated and viable (Fig.5C). Consistent with the observation in zebrafish, only FL *RABEP1* expression rescued the under-agarose migration defect in *RABEP1 knockdown cells.* (Fig. 5D, E; Movie 6).

**Figure 4.**
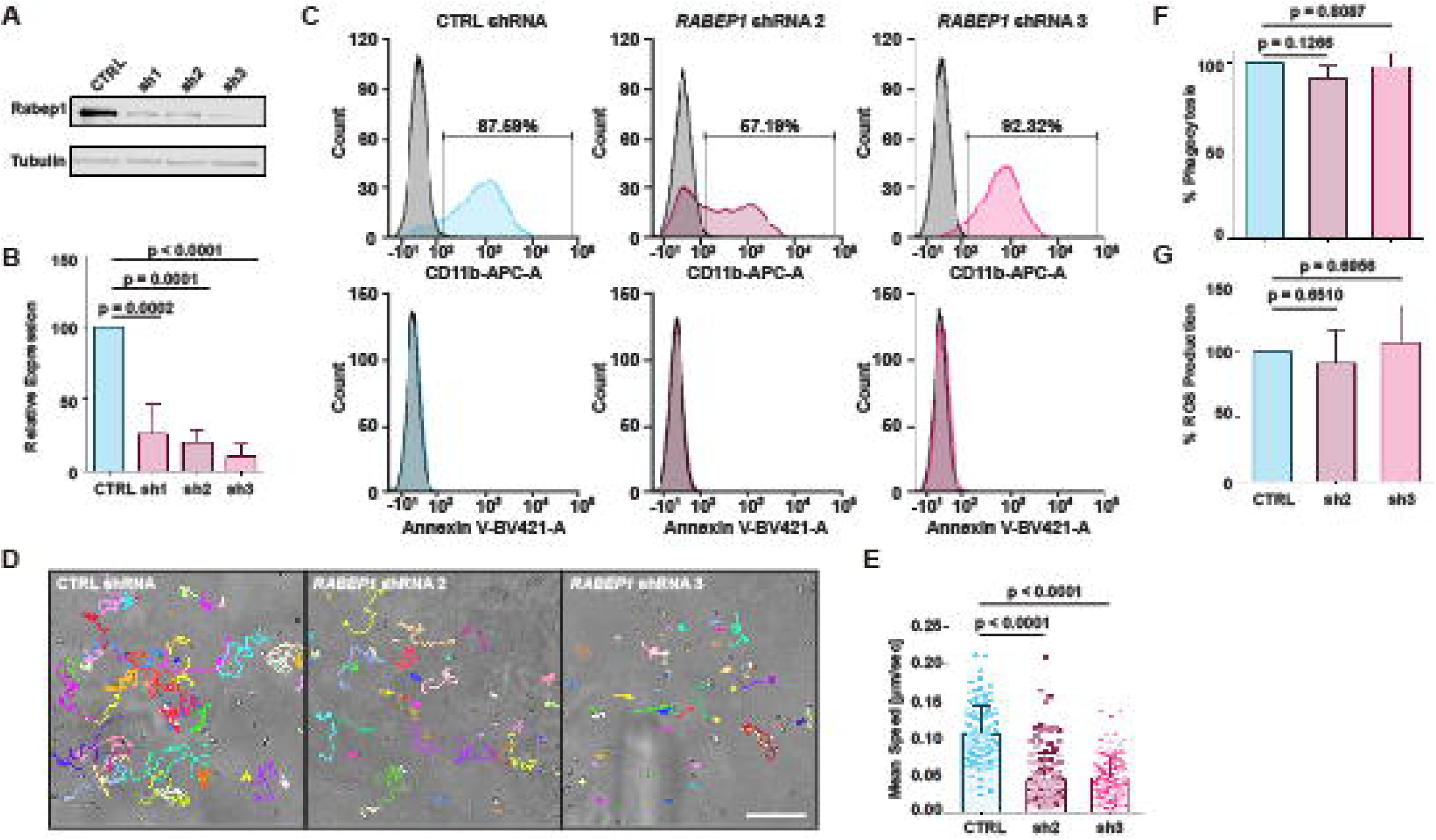
*RABEP1* is essential for motility of HL-60 cells. (A, B) Representative images and quantification for immunoblot of *RABEP1* in control shRNA or *RABEP1* shRNA expressing dHL-60 cells. Tubulin is used as a loading control. Immunoblot was done with 3 biological repeats, and quantification is presented as mean ± SD, using Dunnett’s multiple comparisons test. (C) Representative histogram of CD11b expression and Annexin V staining of control shRNA or *RABEP1* shRNA expressing dHL-60 cells. The positive population of CD11b is gated based on isotype control to show the percentage of HL-60 cells differentiated into neutrophil-like dHL-60 cells. One out of four biological repeats is represented in the image. (D, E) Representative tracks and quantification of the under-agarose random migration of control shRNA or *RABEP1* shRNA expressing dHL-60 cells. (Scale bar, 100μm) The assay was done with 3 biological repeats, each containing 31-58 dHL-60 cells tracked per group. Quantification is presented as mean ± SD, using Dunnett’s multiple comparisons test. (F) Quantification of phagocytosis efficiency of control shRNA or *RABEP1* shRNA expressing dHL-60 cells. The phagocytosis assay using pHrodo^TM^ Green E. coli BioParticles^TM^ was done with 3 biological repeats and is normalized to the geometric mean of control shRNA-expressing dHL-60 cells. Quantification is presented as mean + SD, using Dunnett’s multiple comparisons test. (G) Quantification of ROS production of control shRNA or *RABEP1* shRNA expressing dHL-60 cells. The ROS assay using Amplex^TM^ Red Hydrogen Peroxide/Peroxidase Assay Kit was done with 4 biological repeats and is normalized to the total hydrogen peroxide production of control shRNA expressing dHL-60 cells. Quantification is presented as mean ± SD, using Dunnett’s multiple comparisons test.

**Figure 5.**
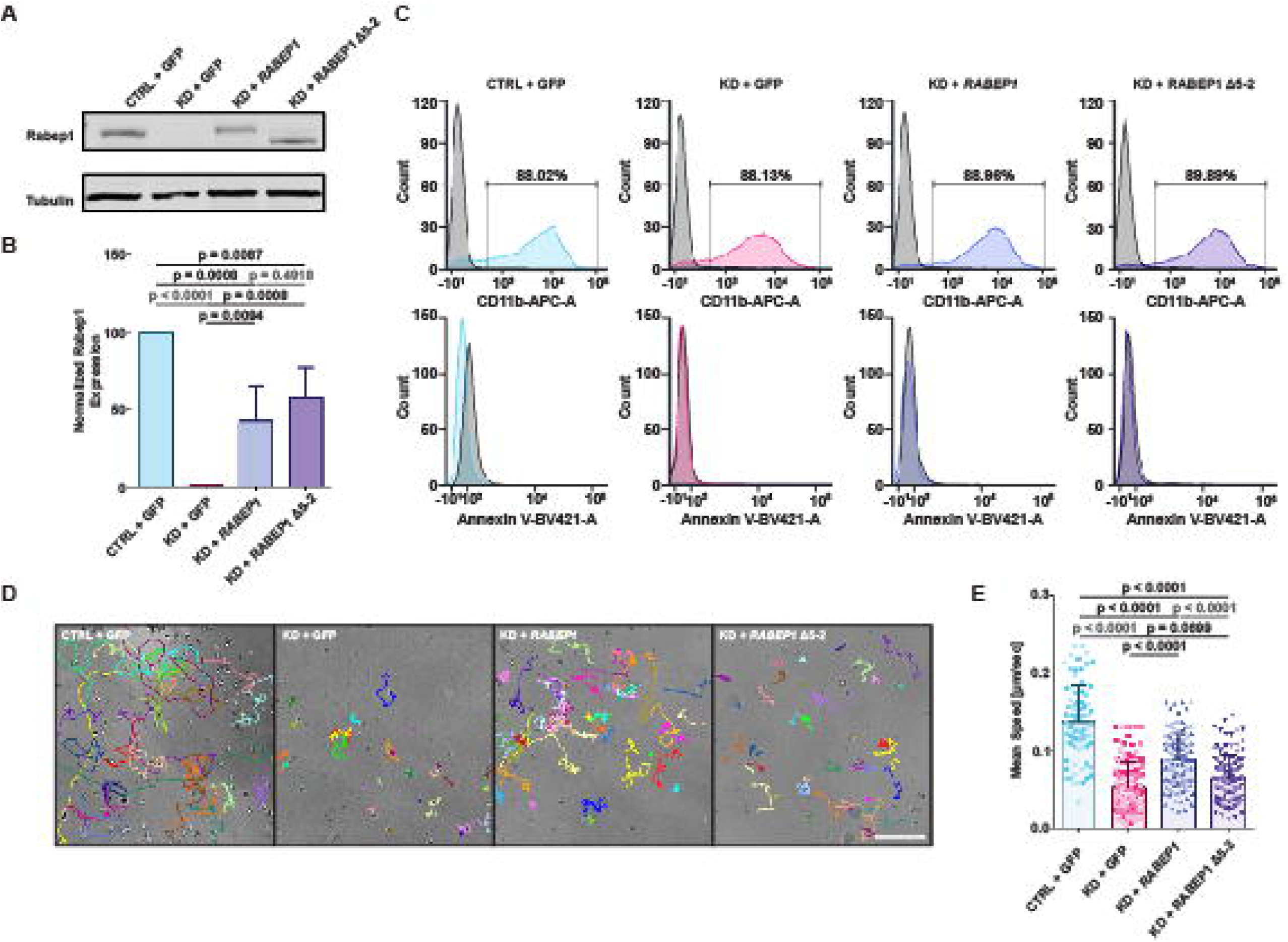
The phenotype of *RABEP1* shRNA knockdown in dHL-60 cells is rescued with FL *RABEP1* but not with Δ5-2 *RABEP1* overexpression. (A, B) Representative images and quantification for immunoblot of *RABEP1* in control shRNA, *RABEP1* shRNA, *RABEP1* shRNA with FL *RABEP1,* or *RABEP1* shRNA with Δ5-2 *RABEP1* expressing dHL-60 cells. Tubulin is used as a loading control. Immunoblot was done with 3 biological repeats, and quantification is presented as mean + SD, using Dunnett’s multiple comparisons test. (C) Representative histogram of CD11b expression and Annexin V staining of control shRNA, *RABEP1* shRNA, *RABEP1* shRNA with FL *RABEP1*, or *RABEP1* shRNA with Δ5-2 *RABEP1* expressing dHL-60 cells. The positive population of CD11b is gated based on isotype control to show the percentage of HL-60 cells differentiated into neutrophil-like dHL-60 cells. One out of four biological repeats is represented in the image. (D, E) Representative images and quantification of tracked under-agarose random migration of control shRNA, *RABEP1* shRNA, *RABEP1* shRNA with FL *RABEP1,* or *RABEP1* shRNA with Δ5-2 *RABEP1* expressing dHL-60 cells. (Scale bar, 100μm) The assay was done with 3 biological repeats, each containing 24-48 dHL-60 cells tracked per group. Quantification is presented as mean ± SD, using Dunnett’s multiple comparisons test.

### *RABEP1* regulates the phosphorylation of P21-activated kinase and actin polymerization

The known function of *RABEP1* is related to early endosome recycling and maturation into late endosomes, where we speculate fast recycling to the plasma membrane via Rab4-GTP recruitment drives downstream pathways, such as Rac activation for actin polymerization, that are essential for precise neutrophil migration (Fig. 6A). To this end, we first labeled the cell membrane with wheat germ agglutinin (WGA) to elucidate endosomal trafficking changes when *RABEP1* is knocked down. In contrast to most cell types with a prominent plasma membrane stain, WGA was absent from the plasma membrane, indicating dynamic endocytosis in dHL-60 cells. WGA accumulated more in the *RABEP1* knockdown cells than in the control, which is also rescued by re-expressing the FL, but not the Δ5-2 *RABEP1* (Fig. 6B, C). This accumulation could also be due to increased internalization, but it supports an endosomal recycling defect. We investigated whether this observed accumulation of endosomes is due to reduced activation of Rab5 stemming from failed recruitment of Rabex5, the Rab5 GEF. Using a Rab5-GTP pull-down assay to measure active Rab5 directly, we did not observe any significant difference between these cell lines (Fig. S2).

**Figure 6.**
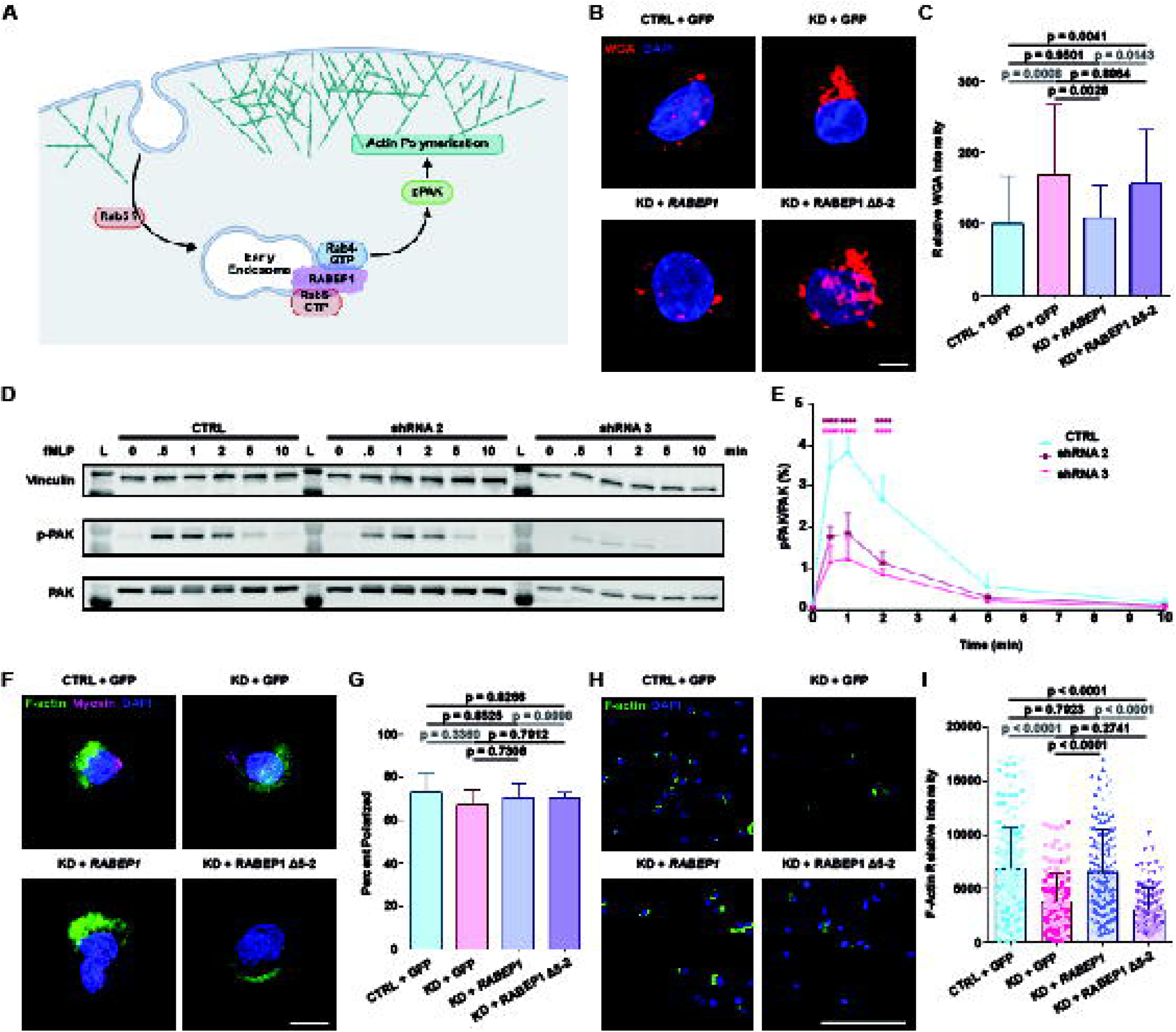
*RABEP1* knockdown dHL-60 cells accumulate endosomes with reduced phosphorylation of PAK and F-actin polymerization. (A) Schematic showing how *RABEP1* mediates endosomal signaling to activate Rac and PAK to form actin filaments at the leading edge. (B, C) Representative images and quantification of WGA IF staining of control shRNA, *RABEP1* shRNA, *RABEP1* shRNA with FL *RABEP1* or *RABEP1* shRNA with Δ5-2 *RABEP1* expressing dHL-60 cells. (Scale bar, 10 μm) The total intensity of each imaged cell was averaged and normalized to control shRNA average intensity values. The assay was done with 2 biological repeats containing 12-33 dHL-60 cells per group. Quantification is presented as mean ± SD, using Dunnett’s multiple comparisons test. (D, E) Representative images and quantification for immunoblot of PAK and pPAK of control shRNA, *RABEP1* shRNA, *RABEP1* shRNA with FL *RABEP1* or *RABEP1* shRNA with Δ5-2 *RABEP1* expressing dHL-60 cells at different points after fMLP stimulation. Vinculin is used as a loading control, and the expression intensity ratio between pPAK and PAK is quantified. Immunoblot was done with 3 biological repeats, and quantification is presented as mean ± SD, using Dunnett’s multiple comparisons test. Significance is presented in asterisks, where 4 asterisks indicate a p-Value of less than 0.0001. (F, G) Representative images and quantification of polarization IF staining of control shRNA, *RABEP1* shRNA, *RABEP1* shRNA with FL *RABEP1* or *RABEP1* shRNA with Δ5-2 *RABEP1* expressing dHL-60 cells. (Scale bar, 10 μm) The ratio of polarized cells to the total number of cells in an image frame was counted. The assay was done with 2 biological repeats, each containing 118-226 dHL-60 cells. Quantification is presented as mean ± SD, using Dunnett’s multiple comparisons test. (H, I) Representative images and quantification of actin intensity measurement of control shRNA, *RABEP1* shRNA, *RABEP1* shRNA with FL *RABEP1* or *RABEP1* shRNA with Δ5-2 *RABEP1* expressing dHL-60 cells. (Scale bar, 100 μm) The assay was done with 2 biological repeats, each containing 31-114 dHL-60 cells quantified per group. Quantification is presented as mean ± SD, using Dunnett’s multiple comparisons test.

It has been shown that endosomal signaling is crucial for Ras-related C3 botulinum toxin substrate 1 (Rac1) activation.^33^ More specifically, chemokine-stimulated CCR7 internalization via endocytosis is essential for activating Rac1. The activation of PAK into phosphorylated PAK (pPAK) by Rac and Cdc42 GTPases is essential in neutrophil migration, where downstream signals of pPAK, such as phosphorylation of myosin light chain (MLC) or activation of actin-binding proteins, are required for proper motility.^34–37^ To determine whether the decreased motility exhibited in *RABEP1* knockdown cell lines is related to this pathway, we measured PAK activation by assessing its phosphorylation using western blot. Indeed, PAK activation, reflected by the pPAK/PAK ratio, is significantly reduced in the two knockdown cell lines across the time points post-fMLP stimulation (Fig. 6D, E). We next investigated whether the cells can polarize to characterize the motility defect in *RABEP1*-deficient cells. All cell lines showed polarized F-actin and myosin light chain (MLC) distribution upon fMLP stimulation, indicating that cell polarization is independent of *RABEP1* (Fig. 6F, G). ^38–39^ However, the total F-actin content is reduced in *RABEP1*-deficient cells, consistent with the function of Rac driving branched actin polymerization at the lamellipodia in the cell front (Fig. 6H, I).

To further elucidate the role of Rab4 and Rab5 in neutrophil migration, we transiently overexpressed dominant negative (DN) Rab4 (S27N) or Rab5 (S34N) in zebrafish neutrophils. Again, both Rab4 DN and Rab5 DN OE neutrophils showed a phenocopy of *rabep1* TSKO neutrophils (Fig.S3A-B; Movie 7).

## DISCUSSION

Here, we report that the miR-190 target gene *RABEP1* is essential for neutrophil migration. We prove that *RABEP1* regulates the recruitment of Rab GTPases to the early endosome for efficient endocytic recycling to the plasma membrane. By visualizing endosome retention and pPAK-F-actin driving the lamellipodia formation, we have identified the mechanism of how *RABEP1* regulates neutrophil motility.

MiR-190 has been implicated in suppressing breast cancer metastasis, where metastatic breast cancer cells exhibited significantly lower levels of *miR-190* expression compared to normal breast cells.^40^ In a different cancer type, hepatocellular carcinoma, *miR-190* facilitates proliferation and metastasis by targeting a tumor suppressor.^41^ Given the expansive target gene pool of *miR-190*, it is being investigated as a potential therapeutic target and biomarker in various diseases, especially cancer.^42–43^ Our initial screening of *miR-190* target gene transient knockout in zebrafish neutrophils revealed several genes that may play a role in cell migration regulation, such as *praf2*, *lats1,* and *rabep1*. Whether these targets also play essential functions in cancer remains to be determined. Furthermore, follow-up studies investigating the other screened *miR-190* target genes, *praf2,* and *lats1*, may reveal novel mechanisms that dictate cell migration.

*RABEP1* is a scaffolding protein that mediates the recruitment of both Rab4-GTP and Rab5-GTP to early endosomes while also acting as an exchange factor of Rab5 through its binding with Rabex5.^44^ Interestingly, it has been reported that Rab5-GTP binds with higher affinity to the C-terminus binding site of Rabaptin-5, whereas Rab4-GTP binds more effectively at the binding site that includes the coil-coiled (CC) 1-2 domain.^30^ The Δ5-2 *RABEP1* lacks a low-affinity binding site for Rab5 GTP and a high-affinity binding site for Rab4 GFP and cannot rescue the *rabep1* TSKO or *RABEP1* shRNA phenotype. We thus speculate that *RABEP1* recruitment of Rab4-GTP to early endosomes is significant for neutrophil migration. Furthermore, Rab5 activation is normal in *RABEP1* knockdown cells, suggesting that *RABEP1* is dispensable for Rab5 activation in dHL-60. It has been shown that the Rabex5 GEF domain is constitutively active but is autoinhibited by its CC domains.^45^ Binding of Rabex5 to Rabaptin-5 allows conformation change to expose the GEF domain and increase GEF activity.^46^ In contrast, others report that Rabex5 GEF activity can be switched on independent of Rabaptin-5 interaction, namely through Rabex5 early endosome targeting (EET) domain, which allows interaction with early endosomes that could relieve autoinhibition of the CCs.^47^ Given the phenocopy of Rab4/5 DN and *rabep1* knockout, our data suggest that *rabep1* binds to Rab5-GTP on early endosomes and then recruits Rab4-GTP to facilitate early endosome recycling, which is essential for neutrophil migration.

Our results are consistent with previous findings where endocytosis is essential for cell migration, albeit not validated in neutrophils before this study. Loss of Sprint, another Rab5 GEF protein, leads to severe migration defects in border cells of *Drosophila* during oogenesis.^48^ Restriction of protein kinase C (PKC) α activation promotes Rab4-dependent recycling of platelet-derived growth factor β receptor (PDGFR) in mouse embryonic fibroblasts (MEFs), which leads to increased chemotaxis of MEFs.^49^ Rabaptin-5 is also found to be a substrate for protein kinase D (PKD) that controls the formation of the Rab4-Rabaptin-5 complex for rapid recycling of α_v_β_3_ integrins, which is needed for driving persistent cell motility and invasion of tumor cells.^50^ Our study elucidates explicitly the importance of Rab4/5-GTP in neutrophil migration in the context of moderating Rac1 activation. Stimulating GPRC and receptor tyrosine kinases leads to the activation of Rac1 via endosomal signaling pathways.^51–52^ Rac1 translocates to the cell surface via recycling endosomes, where it activates the major effector PAK1, which phosphorylates LIM kinase and cortactin to coordinate actin polymerization, and stabilizes newly formed filopodia and lamellipodia.^53^ Therefore, endosomal Rac1 activation plays a central role in cell migration via amplifying the front signal to generate sustained protrusion ^54^

Although there have been a few reports on diseases related to mutations in *RABEP1*, such as increased breast cancer invasion with inhibition of *RABEP1* expression or decreased tumor cell migration when phosphorylation of *RABEP1* is disrupted, the relevance of *RABEP1* in leukocytes has not been established.^55^ Our findings that *RABEP1* is required for neutrophil migration may shed light on the mechanisms governing leukocyte-related diseases, such as acute respiratory distress syndrome (ARDS), rheumatoid arthritis, or sepsis. It may be a potential target for therapeutic strategies.^56–58^

## ACKNOWLEDGEMENTS

We thank our undergraduate students, Caroline Powell and Loahni Hernandez, for cell tracking, counting, and maintaining our zebrafish lines. We thank Dr. David Umulis for providing the LSM880 confocal. Special thanks to Hyein Park for the critical reading of the manuscript and intellectual feedback. We thank the Purdue Imaging Facility for assistance with data collection. The authors gratefully acknowledge the support of the Flow Cytometry and Cell Separation Facility from the Institute for Cancer Research, NIH grant P30 CA023168. This work is supported by NIH-R35GM119787 (to QD).

## AUTHOR CONTRIBUTIONS

DHK: Conceptualization, investigation, data analysis, and writing of manuscript. RS: Investigation, data analysis, and editing of manuscript. CZ, SL, AYH: Investigation, data analysis. JW: supervision. QD: Conceptualization, supervision, funding acquisition, and writing of manuscript.

## DECLARATIONS

### Data Availability Statement

Data supporting the conclusions of this study will be made available by the corresponding author upon reasonable request.

### Competing interests

The authors declare no competing interests.

## ETHICS APPROVAL

All authors read and approve the manuscript.

## Notes

### Competing Interest Statement

The authors have declared no competing interest.

## REFERENCES

1. Nathan, C. Neutrophils and immunity: challenges and opportunities. Nat Rev Immunol 6, 173–182 (2006). 10.1038/nri1785

2. Sadik, C. D., Kim, N. D. & Luster, A. D. Neutrophils cascading their way to inflammation. Trends Immunol 32, 452–460 (2011). 10.1016/j.it.2011.06.008

3. Nauseef, W. M. & Borregaard, N. Neutrophils at work. Nat Immunol 15, 602–611 (2014). 10.1038/ni.2921

4. Witko-Sarsat, V., Rieu, P., Descamps-Latscha, B., Lesavre, P. & Halbwachs-Mecarelli, L. Neutrophils: Molecules, Functions and Pathophysiological Aspects. Lab Invest 80, 617– 653 (2000). 10.1038/labinvest.3780067

5. Segal, A. W. How neutrophils kill microbes. Annu Rev Immunol 23, 197–223 (2005). 10.1146/annurev.immunol.23.021704.115653

6. Fu, X., Liu, H., Huang, G. & Dai, S. The emerging role of neutrophils in autoimmune-associated disorders: effector, predictor, and therapeutic targets. MedComm 2, 402–413 (2021). 10.1002/mco2.85

7. Németh, T., Mócsai, A. & Lowell, C. A. Neutrophils in animal models of autoimmune disease. Semin Immunol 28, 174–186 (2016). 10.1016/j.smim.2016.03.003

8. Chou, R. C. et al. Lipid-Cytokine-Chemokine Cascade Drives Neutrophil Recruitment in a Murine Model of Inflammatory Arthritis. Immunity 33, 266–278 (2010). 10.1016/j.immuni.2010.07.004

9. Dinauer, M. C. Regulation of neutrophil function by Rac GTPases. Curr Opin Hematology 10, 8–15 (2003). 10.1097/00062752-200301000-00003

10. Downey, G. P., Dong, Q., Kruger, J., Dedhar, S. & Cherapanov, V. Regulation of Neutrophil Activation in Acute Lung Injury. Chest 116, 46S–54S (1999). 10.1378/chest.116.suppl_1.46s

11. Christopher, M. J. & Link, D. C. Regulation of neutrophil homeostasis. Curr Opin Hematology 14, 3–8 (2007). 10.1097/MOH.0b013e328011ef5f

12. Ward, J. R. et al. Regulation of Neutrophil Senescence by MicroRNAs. PLoS ONE 6, e15810 (2011). 10.1371/journal.pone.0015810

13. Friedman, R. C., Farh, K. K.-H., Burge, C. B. & Bartel, D. P. Most mammalian mRNAs are conserved targets of microRNAs. Genome Res 19, 92–105 (2009). 10.1101/gr.082701.108

14. Radom-Aizik, S., Zaldivar, F., Oliver, S., Galassetti, P. & Cooper, D. M. Evidence for microRNA involvement in exercise-associated neutrophil gene expression changes. J Appl Physiol 109, 252–261 (2010). 10.1152/japplphysiol.00052.2010

15. Speier, S. et al. Noninvasive high-resolution in vivo imaging of cell biology in the anterior chamber of the mouse eye. Nat Protoc 3, 1278–1286 (2008). 10.1038/nprot.2008.89

16. De La Rosa, I. A. et al. Impaired microRNA processing in neutrophils from rheumatoid arthritis patients confers their pathogenic profile. Modulation by biological therapies. Haematologica 105, 2250–2261 (2020). 10.3324/haematol.2019.230011

17. Lieschke, G. J. & Currie, P. D. Animal models of human disease: zebrafish swim into view. Nat Rev Genet 8, 353–367 (2007). 10.1038/nrg2091

18. Zhou, W. et al. MicroRNA-223 Suppresses the Canonical NF-κB Pathway in Basal Keratinocytes to Dampen Neutrophilic Inflammation. Cell Rep 22, 1810–1823 (2018). 10.1016/j.celrep.2018.01.046

19. Sullivan, C. & Kim, C. H. Zebrafish as a model for infectious disease and immune function. Fish Shellfish Immunol 25, 341–350 (2008). 10.1016/j.fsi.2008.05.005

20. Fan, H.-B., et al. miR-142-3p acts as an essential modulator of neutrophil development in zebrafish. Blood 124, 1320–1330 (2014). 10.1182/blood-2014-03-560839

21. Hsu, A. Y. et al. Phenotypical microRNA screen reveals a noncanonical role of CDK2 in regulating neutrophil migration. Proc Natl Acad Sci USA 116, 18561–18570 (2019). 10.1073/pnas.1902473116

22. Wang, Y. et al. A robust and flexible CRISPR/Cas9-based system for neutrophil-specific gene editing in zebrafish. J Cell Sci 133, jcs240879 (2020). 10.1242/jcs.240879

23. Horiuchi, H. et al. A Novel Rab5 GDP/GTP Exchange Factor Complexed to Rabaptin-5 Links Nucleotide Exchange to Effector Recruitment and Function. Cell 90, 1149–1159 (1997). 10.1016/S0092-8674(00)80357-7

24. Kälin, S., Hirschmann, D. T., Buser, D. P. & Spiess, M. Rabaptin5 is recruited to endosomes by Rab4 and Rabex5 to regulate endosome maturation. Journal of Cell Science jcs.174664 (2015). 10.1242/jcs.174664

25. Van Der Sluijs, P. et al. The small GTP-binding protein rab4 controls an early sorting event on the endocytic pathway. Cell 70, 729–740 (1992). 10.1016/0092-8674(92)80066-W

26. Mohrmann, K., Gerez, L., Oorschot, V., Klumperman, J. & Van Der Sluijs, P. rab4 Function in Membrane Recycling from Early Endosomes Depends on a Membrane to Cytoplasm Cycle. Journal of Biological Chemistry 277, 32029–32035 (2002). 10.1074/jbc.M202470200

27. Dobin A, Davis CA, Schlesinger F, Drenkow J, Zaleski C, Jha S, et al. STAR: Ultrafast Universal RNA-Seq Aligner. Bioinformatics (2013) 29:15–21. 10.1093/bioinformatics/bts635

28. Liao Y, Smyth GK, Shi W. Featurecounts: An Efficient General Purpose Program for Assigning Sequence Reads to Genomic Features. Bioinformatics (2014) 30:923–30. 10.1093/bioinformatics/btt656

29. Huang da W, Sherman BT, Lempicki RA. Systematic and Integrative Analysis of Large Gene Lists Using DAVID Bioinformatics Resources. Nat Protoc (2009) 4:44–57. 10.1038/nprot.2008.211

30. Vitale, G. et al. Distinct Rab-binding domains mediate the interaction of Rabaptin-5 with GTP-bound rab4 and rab5. EMBO J 17, 1941–1951 (1998). 10.1093/emboj/17.7.1941

31. Stenmark, H., Vitale, G., Ullrich, O. & Zerial, M. Rabaptin-5 is a direct effector of the small GTPase Rab5 in endocytic membrane fusion. Cell 83, 423–432 (1995). 10.1016/0092-8674(95)90144-5

32. Bhakta, S. B. et al. Neutrophil-like cells derived from the HL-60 cell-line as a genetically-tractable model for neutrophil degranulation. PLoS ONE 19, e0297758 (2024). 10.1371/journal.pone.0297758

33. Hahn, H. et al. Endosomal chemokine receptor signalosomes regulate central mechanisms underlying cell migration. eLife (2025). 10.7554/eLife.08384

34. Chan, P. M., Lim, L. & Manser, E. PAK Is Regulated by PI3K, PIX, CDC42, and PP2Cα and Mediates Focal Adhesion Turnover in the Hyperosmotic Stress-induced p38 Pathway. Journal of Biological Chemistry 283, 24949–24961 (2008). 10.1074/jbc.M804917200

35. Chu, J. et al. Biphasic Regulation of Myosin Light Chain Phosphorylation by p21-activated Kinase Modulates Intestinal Smooth Muscle Contractility. Journal of Biological Chemistry 288, 1200–1213 (2013). 10.1074/jbc.M112.439267

36. Arber, S. et al. Regulation of actin dynamics through phosphorylation of cofilin by LIM-kinase. Nature 393, 805–809 (1998). 10.1038/31642

37. Yang, N. et al. Cofilin phosphorylation by LIM-kinase 1 and its role in Rac-mediated actin reorganization. Nature 393, 809–812 (1998). 10.1038/31511

38. Gardel, M. L., Schneider, I. C., Aratyn-Schaus, Y. & Waterman, C. M. Mechanical Integration of Actin and Adhesion Dynamics in Cell Migration. Annu. Rev. Cell Dev. Biol. 26, 315–333 (2010). 10.1146/annurev-cellbio-100109-104028

39. Byrne, K. M. et al. Bistability in the Rac1, PAK, and RhoA Signaling Network Drives Actin Cytoskeleton Dynamics and Cell Motility Switches. Cell Systems 2, 38–48 (2016). 10.1016/j.cels.2016.06.012

40. Yu, Y. et al. miR-190 suppresses breast cancer metastasis by regulation of TGF-β-induced epithelial–mesenchymal transition. Mol Cancer 17, 70 (2018). 10.1186/s12885-018-0372-4

41. Xiong, Y. et al. miR-190 promotes HCC proliferation and metastasis by targeting PHLPP1. Experimental Cell Research 371, 185–195 (2018). 10.1016/j.yexcr.2018.07.004

42. Yu, Y. et al. miR-190 enhances endocrine therapy sensitivity by regulating SOX9 expression in breast cancer. J Exp Clin Cancer Res 38, 22 (2019). 10.1186/s13046-019-0997-7

43. De Lella Ezcurra, A. L., et al. miR-190 Enhances HIF-Dependent Responses to Hypoxia in Drosophila by Inhibiting the Prolyl-4-hydroxylase Fatiga. PLoS Genet 12, e1006073 (2016). 10.1371/journal.pgen.1006073

44. Mattera, R., Tsai, Y. C., Weissman, A. M. & Bonifacino, J. S. The Rab5 Guanine Nucleotide Exchange Factor Rabex-5 Binds Ubiquitin (Ub) and Functions as a Ub Ligase through an Atypical Ub-interacting Motif and a Zinc Finger Domain. Journal of Biological Chemistry 281, 6874–6883 (2006). 10.1074/jbc.M510595200

45. Lauer, J. et al. Auto-regulation of Rab5 GEF activity in Rabex5 by allosteric structural changes, catalytic core dynamics and ubiquitin binding. eLife 8, e46302 (2019). 10.7554/eLife.46302

46. Zhang, Z. et al. Molecular mechanism for Rabex-5 GEF activation by Rabaptin-5. eLife 3, e02687 (2014). 10.7554/eLife.02687

47. Zhu, H. et al. Rabaptin-5-independent Membrane Targeting and Rab5 Activation by Rabex-5 in the Cell. Mol Biol Cell 18, 4119–4128 (2007). 10.1091/mbc.E07-02-0111

48. Jékely, G., Sung, H.-H., Luque, C. M. & Rørth, P. Regulators of Endocytosis Maintain Localized Receptor Tyrosine Kinase Signaling in Guided Migration. Developmental Cell 9, 197–207 (2005). 10.1016/j.devcel.2005.06.014

49. Hellberg, C., Schmees, C., Karlsson, S., Åhgren, A. & Heldin, C.-H. Activation of Protein Kinase C Is Necessary for Sorting the PDGF-Receptor to Rab4a-dependent Recycling. Mol Biol Cell 20, 2856–2863 (2009). 10.1091/mbc.E08-07-0764

50. Christoforides, C., Rainero, E., Brown, K. K., Norman, J. C. & Toker, A. PKD Controls αvβ3 Integrin Recycling and Tumor Cell Invasive Migration through Its Substrate Rabaptin-5. Developmental Cell 23, 560–572 (2012). 10.1016/j.devcel.2012.07.009

51. Laufer, Julia M., Mark A. Hauser, Ilona Kindinger, Vladimir Purvanov, Andreas Pauli, and Daniel F. Legler. Chemokine Receptor CCR7 Triggers an Endomembrane Signaling Complex for Spatial Rac Activation. Cell Reports 29, no. 4 (October 2019): 995–1009.e6. 10.1016/j.celrep.2019.09.031.

52. Palamidessi, Andrea, Emanuela Frittoli, Massimiliano Garré, Mario Faretta, Marina Mione, Ilaria Testa, Alberto Diaspro, Letizia Lanzetti, Giorgio Scita, and Pier Paolo Di Fiore. Endocytic Trafficking of Rac Is Required for the Spatial Restriction of Signaling in Cell Migration. Cell 134, no. 1 (July 2008): 135–47. 10.1016/j.cell.2008.05.034.

53. Schiefermeier, Natalia, David Teis, and Lukas A Huber. Endosomal Signaling and Cell Migration. Current Opinion in Cell Biology 23, no. 5 (October 2011): 615–20. 10.1016/j.ceb.2011.04.001.

54. Tebar, Francesc, Carlos Enrich, Carles Rentero, and Thomas Grewal. GTPases Rac1 and Ras Signaling from Endosomes. In Endocytosis and Signaling, edited by Christophe Lamaze and Ian Prior, 57:65–105. Progress in Molecular and Subcellular Biology. Cham: Springer International Publishing, 2018. 10.1007/978-3-319-96704-2_3.

55. Wei, F., Cao, C., Xu, X. & Wang, J. Diverse functions of miR-373 in cancer. J Transl Med 13, 162 (2015). 10.1186/s12967-015-0500-x

56. Cortjens, B., Van Woensel, J. B. M. & Bem, R. A. Neutrophil Extracellular Traps in Respiratory Disease: guided anti-microbial traps or toxic webs? Paediatric Respiratory Reviews 21, 54–61 (2017). 10.1016/j.prrv.2016.11.006

57. Villanueva, E. et al. Netting Neutrophils Induce Endothelial Damage, Infiltrate Tissues, and Expose Immunostimulatory Molecules in Systemic Lupus Erythematosus. The Journal of Immunology 187, 538–552 (2011). 10.4049/jimmunol.1100326

58. Herrero-Cervera, A., Soehnlein, O. & Kenne, E. Neutrophils in chronic inflammatory diseases. Cell Mol Immunol 19, 177–191 (2022). 10.1038/s41423-021-00747-7

