## Supplemental figure and movie legend for "RABEP1 amplifies front signaling in neutrophil migration"

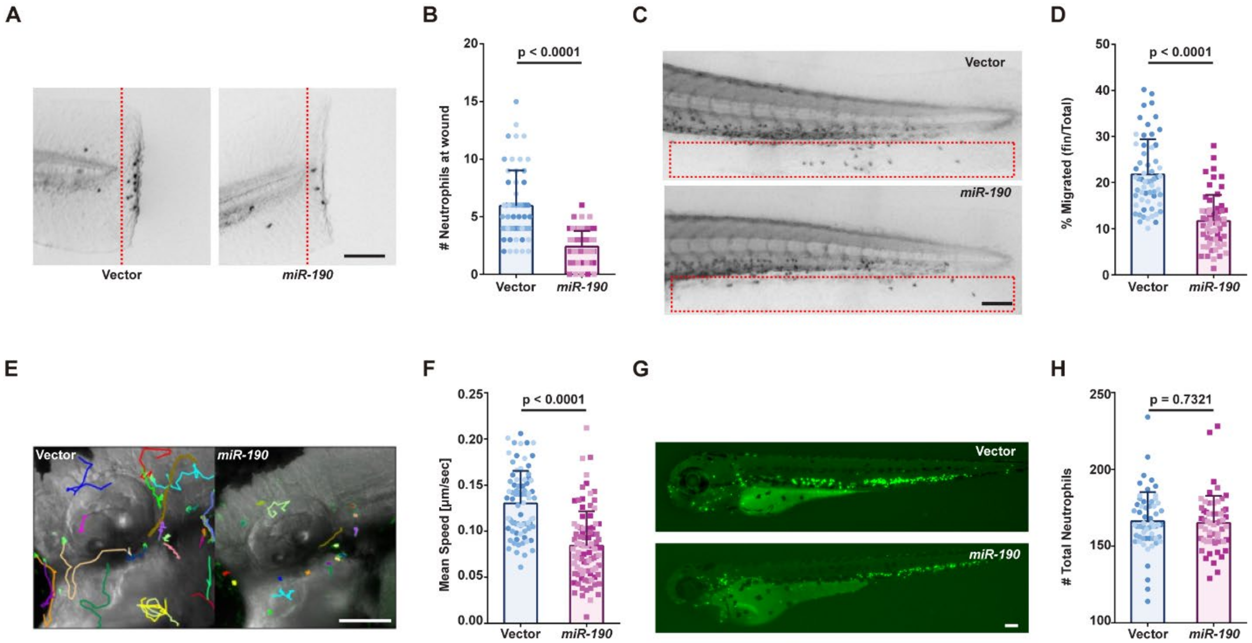


**Figure S1. Impaired neutrophil motility in a second *miR-190* OE founder fishline.** (A, B) Representative images and quantification of neutrophil recruitment to the tail wound site in vector and miR-190 OE larvae from the second founder, 1 hour post-wounding. (Scale bar, 100 µm) The assay was done with 3 biological repeats, each containing 17-22 fish per group. Quantification is presented as mean ± SD, using the Mann-Whitney test. (C, D) Representative images and quantification of neutrophil chemotaxis percentage from CHT to caudal fin in vector and *miR-190* OE larvae from the second founder with 15min LTB4 exposure. (Scale bar, 100 µm) The assay was done with 3 biological repeats, each containing 20 fish per group. Quantification is presented as mean ± SD, using the Mann-Whitney test. (E, F) Representative images and quantification of live neutrophil motility imaging in vector and *miR-190* OE larvae from the second founder for 30 min. (Scale bar, 100µm) The assay was done with 3 biological repeats, each containing 6-39 neutrophils tracked from 1-3 fish per group. Quantification is presented as mean ± SD, using the Mann-Whitney test. (G, H) Representative GFP images and Sudan black staining quantification of whole-body neutrophil count in vector and *miR-190* OE larvae from the second founder. (Scale bar, 100µm) The assay was done with 3 biological repeats, each containing 20 fish per group. Quantification is presented as mean + SD, using the Mann-Whitney test. All vector data is the same as Figure 1, and experiments were done together.

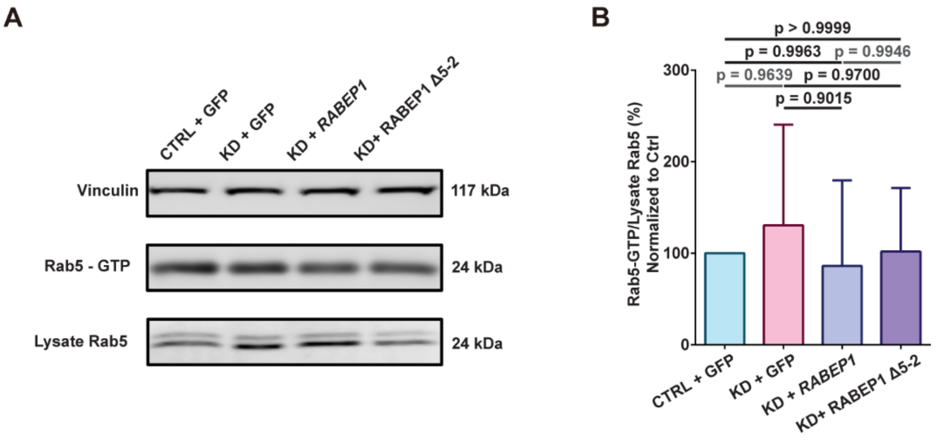


**Figure S2.** ***RABEP1*** **knockdown does not change Rab5 activation.** (A, B) Representative images and quantification for immunoblot of Rab5-GTP in control shRNA, *RABEP1* shRNA, *RABEP1* shRNA with FL *RABEP1,* or *RABEP1* shRNA with Δ5-2 *RABEP1* expressing dHL-60 cells. Vinculin is used as a loading control, and the expression intensity ratio between precipitated Rab5-GTP and lysate Rab5 is quantified. Immunoblot was done with 3 biological repeats, and quantification is presented as mean ± SD, using Dunnett's multiple comparisons test.

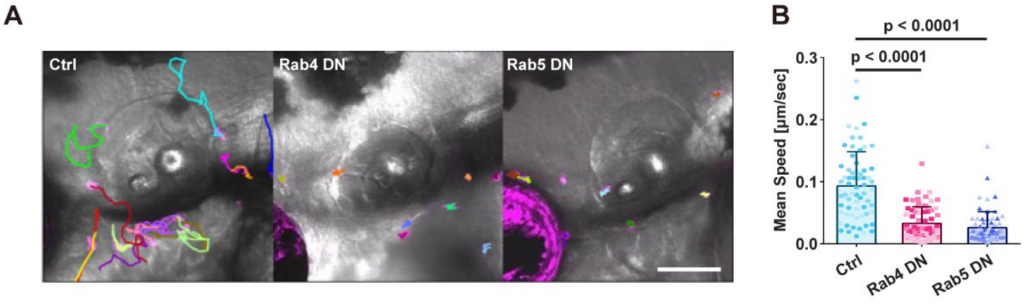


**Figure S3. Rab4 DN and Rab5 DN reduced neutrophil motility in zebrafish.** (A, B) Representative images and quantification of live neutrophil motility imaging in Rab4 DN OE, Rab5 DN OE, and Rab11 DN OE larvae. (Scale bar, 100 µm.) The assay was done with 3 biological repeats, each containing 17-29 neutrophils tracked from 1-5 fish per group. Quantification is presented as mean ± SD, using Dunnett's multiple comparisons test.

Movie legend

**Movie 1. Tracked movies of neutrophil motility in the head mesenchyme of vector and *miR-190* from the first F0 zebrafish.** The movie shows the migration of neutrophils in 3-dpf *Tg(LyzC-miR-190-dendra2)$^{pu38}$* F0 #1 and *Tg(LyzC-vector-dendra2)$^{pu7}$*. (Scale bar, 100 μm)

**Movie 2. Tracked movies of neutrophil motility in the head mesenchyme of vector and *miR-190* from the second F0 zebrafish.** The movie shows the migration of neutrophils in 3 dpf *Tg(LyzC-miR-190-dendra2)$^{pu38}$* F0 #2 and *Tg(LyzC-vector-dendra2)$^{pu7}$*. (Scale bar, 100 μm)

**Movie 3. Tracked movies of neutrophil motility in the head mesenchyme of control TSKO and *rabep1* TSKO zebrafish.** The movie shows the migration of neutrophils in 3 dpf *Tg(LyzC: ctrl sgRNAs, LyzC: Cas9, Cry: RFP)* and *Tg(LyzC: rabep1 sgRNAs, LyzC: Cas9, Cry: RFP).* (Scale bar, 100 μm)

**Movie 4. Tracked movies of neutrophil motility in the head mesenchyme of ctrl rescue, FL *RABEP1* rescue, and Δ5-2 *RABEP1* rescue zebrafish.** The movie shows the migration of neutrophils in 3-dpf larvae from crossing *Tg(LyzC: rabep1 sgRNAs, LyzC: Cas9, Cry: RFP)* with *Tg(LyzC:RFP-Caax)$^{pu54,}$* *Tg(LyzC:FL RABEP1-RFP-Caax)$^{pu53}$* and *Tg(LyzC: Δ5-2 RABEP1-RFP-Caax)$^{pu55}$*. (Scale bar, 100 μm)

**Movie 5. Tracked movies of ctrl shRNA dHL-60 cells and *RABEP1* shRNA dHL-60 cells.** The movie shows the under-agarose migration of ctrl shRNA dHL-60 cells and *RABEP1* shRNA dHL-60 cells. (Scale bar, 100 μm)

**Movie 6. Tracked movies of Ctrl rescue dHL-60 cells, FL RABEP1 rescue dHL-60 cells, and Δ5-2 *RABEP1* rescue dHL-60 cells.** The movie shows the under-agarose migration of control rescue dHL-60 cells, FL *RABEP1* rescue dHL-60 cells, and Δ5-2 *RABEP1* rescue dHL-60 cells. (Scale bar, 100 μm)

**Movie 7. Tracked movies of neutrophil motility in the head mesenchyme of Rab4 DN OE and Rab5 DN OE zebrafish.** The movie shows the migration of neutrophils in 3-dpf Rab4 DN OE and Rab5 DN OE zebrafish lines. (Scale bar, 100 μm)
